# Adaptive problem solving in the primate frontal cortex

**DOI:** 10.64898/2026.05.04.722785

**Authors:** Mahdi Ramadan, Adam Gosztolai, Mehrdad Jazayeri

## Abstract

Humans solve problems adaptively by selecting strategies suited to the situation. For example, when missing the bus to an appointment, we may wait for the next bus, call a taxi, cancel, or reschedule depending on the circumstances. Yet the neural and computational principles that support such flexible problem solving remain poorly understood. To address this question, we designed a moderately complex decision task for monkeys that allows multiple plausible solution strategies. Animals learned the task rapidly, generalized to novel maze geometries, and their choices were inconsistent with any single fixed strategy. We then recorded large-scale neural activity from the frontal cortex and found that population dynamics varied systematically with maze geometry. Neural responses clustered into two distinct dynamical regimes with separable initial states, consistent with hierarchical and sequential strategies. A decoder trained on population activity revealed time-resolved decision dynamics that aligned with these regimes, and an unsupervised latent-space analysis provided convergent evidence that strategy use varied across trials. A behavioral model grounded in neurally inferred strategies accounted for choices better than fixed-strategy alternatives and captured trial-by-trial variability. Together, these results provide a neural and computational account of how the brain selects and implements distinct strategies during adaptive problem solving.

## Introduction

A core component of human intelligence is the capacity to assess problems and devise suitable strategies to solve them. For instance, what we do when our phone goes missing depends on the situation: at home, we might ask others; at work, we might retrace our steps; in a store, we might check with the cashier; and on vacation, we might attempt to track it online (Fig. 1a). A fundamental open question in cognitive, computational and systems neuroscience is how the nervous system selects and implements adaptive strategies for problem solving.

**Figure 1.**
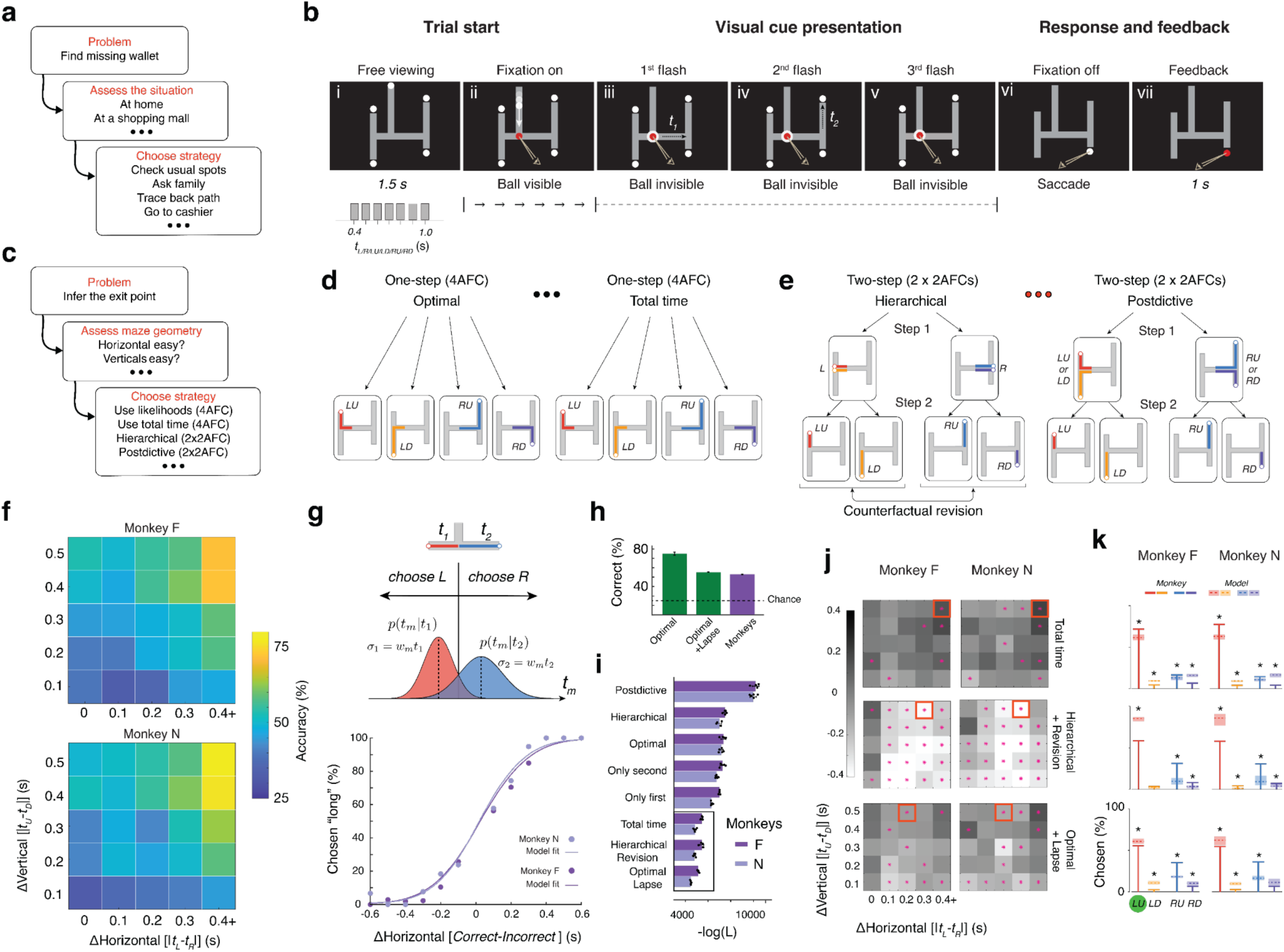
Task, strategies, and behavior. (a) A real-world example of problem solving highlighting the need for (i) assessing the situation and (ii) selecting a suitable strategy. (b) Task. (i) An H-shaped maze and four possible choices are presented. (ii) After monkeys fixate a central red circle, a visible ball begins to move downward toward the maze with constant speed. (iii) A flash signals when the ball turns leftward or rightward. From this point onward, the ball is invisible. (iv) Another flash signals when the ball reaches the vertical segment and turns upward or downward (*t_1_*: interval between the first and second flashes). (v) A third flash signals when the ball reaches an endpoint (*t_2_*: interval between the second and third flashes). Note that all flashes are presented centrally around the fixation point and not where the ball is. (vi) The fixation spot disappears and monkeys report the inferred exit point with a saccade. (vii) Monkeys receive binary visual feedback only at the location of their choice and a juice reward for a correct answer. The distribution from which all arm lengths (2 horizontal and 4 vertical) were sampled are shown below panel (i). The maze segment lengths are measured in units of time and denoted by *t_L_* (left), *t_R_* (right), *t_LU_* (left-up), *t_LD_* (left-down), *t_RU_* (right-up), *t_RD_* (right-down). (c) Computational steps for solving the H-maze task: (i) Monkeys may assess the maze geometry (e.g., the discriminability of horizontal and vertical arms) and then (ii) choose among various 4AFC (choosing between the 4 exits) and 2AFCs (one for the horizontal segments and another for the vertical segments) strategies. (d) Two example 4AFC strategies. Left: Optimal. This model computes the joint likelihood of the horizontal and vertical arms (colored arms) leading to each exit point (four panels) and chooses the most likely path based on *t_1_* and *t_2_*. Right: Total time. This model computes the likelihood of the total path leading to each exit point (four panels) and chooses the most likely one based on *t_1_+t_2_*. Note that the decisions are made based on noisy internal estimates of *t_1_* and *t_2_*, and not their objective values (see Methods). (e) Two example 2AFC strategies. Left: Hierarchical. This model first chooses left versus right based on *t_1_* (top two panels) and then between the connected vertical arms based on *t_2_* (two bottom pairs of panels). The arrow denoted “Counterfactual revision” shows a variant in which the model could revise its left/right decision if the likelihood of the vertical arms under consideration is below a threshold (see Methods for details). Right: Postdictive. This model behaves similarly to the hierarchical model with the difference that it chooses left versus right postdictively based on both *t_1_* and *t_2_*. (f) Task performance. Percentage of correct responses as a function of absolute length difference between the two horizontal arms (*|t_L_-t_R_|*) and the two vertical arms on the side chosen by monkeys (*|t_LU_-t_RD_|* for left and *|t_RU_-t_RD_|* for right). In the plot, we label the ordinate as *|t_U_-t_D_|*, which combines both left and right responses. (g) Top: T-maze task, which is identical to the H-maze without the vertical arms. Middle: The model. The animal uses its noisy estimate of the sample time interval (*t_1_* and *t_2_*) denoted *t_m_*, to choose the more likely alternative. We assumed *t_m_* is subject to scalar variability and modeled the conditional probability of *t_m_* on the sample interval as a Gaussian distribution whose standard deviation is proportional to its mean with the constant of proportionality denoted *w_m_*. We fit this model to two of the animals’ choices to estimate *w_m_*. Bottom: Percent chosen “long” as a function of the difference between the correct and incorrect segments for the two animals (circles) as well as the corresponding model fits (lines). (h) Suboptimal performance. Animals’ performance was lower than predicted by the model instantiating an optimal strategy (see Methods). The performance could be matched to an optimal model with a fitted lapse rate. (i) Negative log-likelihood of different fixed strategy models shown for the two animals, separately (see Methods). The box at the bottom highlights the top three model candidates. (j) Log performance ratio of animals to the three top models highlighted in panel i separated for different maze geometries. Asterisks (*) indicates statistical significance of difference between animals and models (two-sided t-tests, α = 0.05). Red squares are the geometries with the largest discrepancy. Positive values indicated better monkey performance. (k) Comparison of choice distributions between animals and the three top fixed-strategy models (box in panel i) for specific maze geometries (red squares in panel j). The choice probabilities for the four exits (*LU*, *LD*, *RU*, *RD*) are shown as colored lines (animals: solid; models: dashed). The shaded regions are the bounding boxes over 5 model bootstraps. To facilitate visualization, all mazes were rotated and/or reflected so that *LU* represents the correct exit (green circle). Asterisks (*) denote statistically significant differences (two-tailed t-tests, α = 0.05) between monkey and model.

Cognitive scientists have long studied problem solving in humans under the rubrics of mental algorithms, heuristics, and reasoning ^1–6^. Computational accounts have proposed mechanisms such as contextual inference and probabilistic reasoning as key components of this process ^7–16^. In parallel, neuroimaging studies have identified candidate anatomical substrates for these computations ^17–24^. Problem solving is also a major target of research in modern machine learning aimed at engineering general artificial intelligence ^25–31^. However, despite these advances, the neural mechanisms that enable flexible selection and deployment of strategies remain poorly understood.

Animal studies, where neural mechanisms can be examined with high precision, have traditionally focused on relatively simple, one-step decision tasks ^32–60^. These paradigms have enabled detailed characterization of optimal and near-optimal decision policies. More recent work has extended this approach to tasks that feature context-dependent computations, where context is either provided explicitly ^55,61–66^ or inferred from experience ^53,67–71^. However, since these paradigms do not motivate or require animals to endogenously contextualize the problem, the representational principles and dynamical motifs that support strategic problem solving remain poorly understood.

We developed a moderately complex temporal decision-making task, the H-maze, that admits multiple plausible task representations and solution strategies. Crucially, the task allows animals either to rely on a single fixed strategy across all trials or to flexibly deploy different strategies depending on the specific geometry. Behavioral analyses showed that animals solved the task accurately and generalized to novel maze configurations, yet their choice patterns were inconsistent with any fixed-strategy solution.

To identify the strategies animals used and uncover the underlying neural computations, we performed large-scale electrophysiological recordings in the dorsomedial frontal cortex (DMFC), a region implicated in temporal decision-making ^62–65,72–80^. Neural activity organized into geometry-dependent dynamical regimes, consistent with the use of strategies adapted to task demands. These dynamics were associated with distinct initial states that predicted the ensuing regime on a trial-by-trial basis. Finally, animals’ choice behavior across trials and maze geometries was best captured by a latent behavioral model in which strategy was inferred from neural activity. Together, these results provide a neural basis for the selection and implementation of adaptive computational strategies during problem solving.

## Results

### H-maze task

In each trial, a ball moved at constant speed through an H-shaped maze whose arm lengths varied across trials (Fig. 1b; Supplementary Video 1). The ball was visible as it entered from above the central horizontal segment. From that point onward, it became invisible, and its trajectory had to be inferred from three precisely timed flashes presented at the fixation point. The first flash signaled entry into the horizontal arm, the second signaled entry into a vertical arm, and the third signaled arrival at an exit. Shortly after the ball stopped, the fixation point disappeared, at which point the monkey had to report the exit location with a saccade.

Because the ball moved at constant speed, solving the task required mapping the relative timing of flashes onto the lengths of the maze segments. We define *t_1_* as the interval between the first and second flashes (horizontal travel time) and *t_2_* as the interval between the second and third flashes (vertical travel time). We express arm lengths in units of time: *t_L_* (left) and *t_R_* (right), *t_LU_* (left-up), *t_LD_* (left-down), *t_RU_* (right-up), and *t_RD_* (right-down). For the initial behavioral experiments, each arm was sampled independently from the set {0.4, 0.5, 0.6, 0.7, 0.8, 0.9, 1.0 s} (Fig. 1b, inset), yielding 7^6^ unique maze geometries. This large stimulus set ensured that animals could not rely on memorized stimulus–response mappings to solve the task.

### A rich hypothesis space for task representations and strategies

The H-maze task admits multiple task representations and solution strategies (Fig. 1c). At the representational level, animals could treat the task either as a four-alternative forced choice (4AFC), selecting directly among the four exits, or as two sequential two-alternative forced choice decisions (2×2AFC): first left versus right, then up versus down. Each representation, in turn, supports multiple strategies.

Under the 4AFC representation, an optimal strategy is to compute the joint likelihood of each full path and select the most likely one given *t_1_* and *t_2_* (Fig. 1d, left). Simpler heuristics are also plausible, such as matching the total elapsed time *t_1_* + *t_2_* to total path length (Fig. 1d, right), or relying only on *t_2_* to select among vertical arms.

The 2×2AFC representation is also consistent with multiple strategies. A hierarchical strategy first uses *t_1_* to choose between left and right and then uses *t_2_* to choose between the corresponding vertical arms (Fig. 1e, left). This strategy can be augmented by counterfactual revision: if the vertical arms associated with the initial horizontal choice are inconsistent with *t_2_* beyond an acceptable threshold, the initial decision can be revised. Alternatively, a postdictive strategy combines *t_1_* and *t_2_* to infer left versus right and then uses *t_2_* to resolve the vertical decision (Fig. 1e, right).

Crucially, the task also allows for adaptive strategies based on geometry. For example, when the horizontal arms are easily discriminable, a hierarchical strategy is advantageous, whereas when they are similar, strategies that rely on *t_2_* are more effective. In what follows, we first show that animals perform the task accurately, then demonstrate that their behavior cannot be explained by any fixed strategy and finally use neural recordings to identify and characterize the strategies deployed on a trial-by-trial basis.

### Monkeys solve H-maze flexibly

We first trained monkeys on a version of the task in which the ball remained visible throughout its trajectory (Video S1). Animals rapidly learned to report the correct exit and reached near-ceiling performance on trained maze geometries (Monkey F: 99.08 ± 0.85%; Monkey N: 100%). They also generalized to untrained geometries (Fig. S1a), indicating that they had acquired a general understanding of the task structure.

We then trained animals on the main task in which the ball was invisible (Video S1). After performance stabilized, we tested whether animals used both intervals to guide their decisions. A multivariate regression analysis revealed that choices depended on both the discriminability of the horizontal arms (ΔH) and the discriminability of the vertical arms on the chosen side (ΔV) (Fig. 1f, Monkey F: *β*_ΔH_[CI_ΔH_]=0.166[0.147-0.184], *β*_ΔV_[CI_ΔV_]=0.191[0.170-0.212], *p*=5.343e-67/9.683e-73; Monkey N: *β*_ΔH_[CI_ΔH_]=0.199[0.179-0.219], *β*_ΔV_[CI_ΔV_]=0.213[0.191-0.235], *p*=2.193e-82/5.411e-80; *H_0_*: *β*=0). These results indicate that, across mazes, animals used both *t_1_* and *t_2_* to solve the task.

A hallmark of adaptive problem solving is the ability to flexibly generalize and evaluate alternatives across contexts. We therefore tested animals on multiple forms of generalization. In a parametric generalization test, animals were presented with mazes containing previously unseen arm lengths and performed accurately (Fig. S1b). In a more stringent structural generalization, we confronted animals with novel maze configurations with different numbers of turns and exits, which they also solved within a single session (Fig. S1c). Beyond generalization, analysis of post-feedback eye movements revealed that animals evaluated counterfactual outcomes in a manner consistent with their probabilities, indicating that they tracked unchosen alternatives and their expected outcomes (Fig. S1d). Together, these results provide convergent evidence that animals approached the task with a high degree of cognitive flexibility.

Together, these results indicate that animals acquired a flexible solution to the task that goes beyond statistical learning.

### Behavioral responses are not consistent with a fixed strategy

To test whether animals’ choices could be explained by a single fixed strategy, we compared their behavior to predictions from a broad set of candidate models. To ensure a stringent comparison, we adopted an Occam’s razor approach: we first estimated each animal’s timing variability using a separate two-alternative forced choice (2AFC) T-maze task (Fig. 1g), and then used this independently estimated parameter to generate a priori predictions for each fixed-strategy model without refitting each animal’s timing variability.

In the T-maze task, animals discriminated between two intervals corresponding to the horizontal arm lengths. Behavior was well captured by a model with scalar timing variability ^81^, parameterized by a Weber fraction *w_m_*. Fitting this model yielded accurate estimates of timing noise for each animal (Monkey F: 0.165; Monkey N: 0.153; Fig. 1g), consistent with the scalar property of timing.

We then used these independently estimated parameters to construct predictive models implementing different fixed strategies and compared their choice distributions to those of the animals. None of the fixed-strategy models accounted for behavior accurately; all exhibited systematic deviations across maze geometries. The optimal model outperformed the animals (Fig. 1h), consistent with prior findings in humans ^82^. Models instantiating hierarchical and postdictive strategies were similarly deficient (Fig. 1i).

We also tested models implementing a range of heuristic strategies, including those that relied only on the first interval, only on the second interval, or on the total elapsed time. All such models failed to capture animals’ behavior, exhibiting systematic deviations across maze geometries (Fig. 1j,k; Fig. S2). These deviations were not random, but followed consistent patterns that differed from model predictions. For example, an optimal model augmented with a lapse rate to match overall performance (Fig. 1h) systematically overpredicted leftward choices in mazes with intermediate left–right discriminability (Fig. 1k, bottom). Notably, these discrepancies were consistent across animals, indicating structured biases rather than idiosyncratic noise. Together, these results show that animals’ behavior cannot be explained by any single fixed strategy, and instead, suggest that animals adapt their strategy across maze geometries.

### Structured neural activity in the frontal cortex

To gain insight into the strategies animals used, we examined neural activity in a region of DMFC (Fig. 2a, S3) that supports interval timing, temporal decision-making, and tracking invisible stimuli ^62–65,72–80,83^. We used Neuropixels probes to record simultaneously from large populations of neurons, enabling us to characterize population dynamics on a trial-by-trial basis.

**Figure 2.**
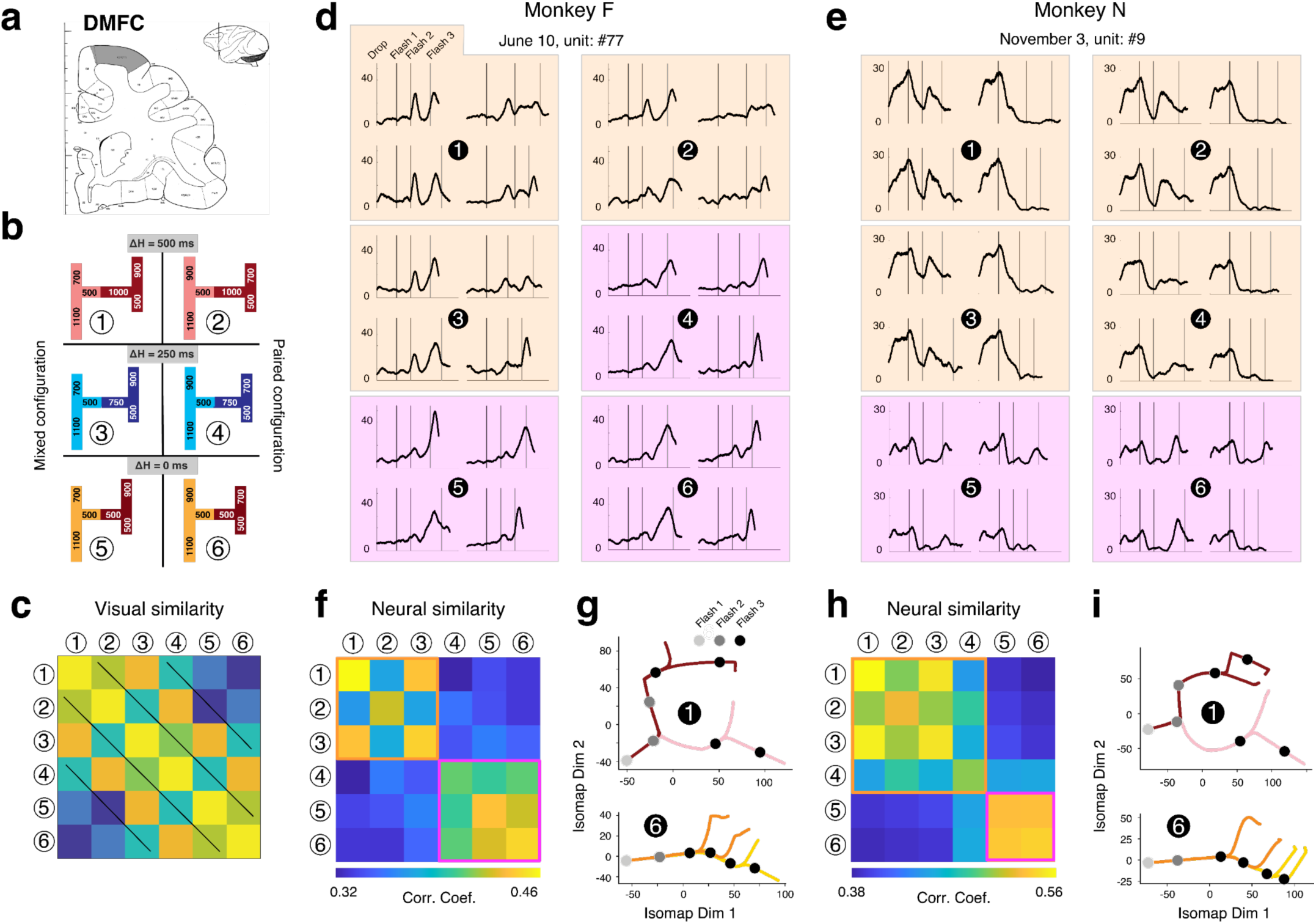
Single-neuron categorical geometry-dependent response dynamics. **(a)** Recording sites. The coronal section is aligned to the vertical line in the inset, with DMFC highlighted in gray. **(b)** The 6 maze geometries used in the physiology experiment. Rows are ordered according to the difference between the two horizontal arms. Left column: “Paired” configuration. In this configuration, the vertical arms on the left are both longer than the two on the right. Right column: “Mixed” configuration. In this configuration, the lengths of vertical arms are mixed between the left and right side of the maze. **(c)** The visual similarity between mazes. We encoded each maze by a 6-vector containing its 2 horizontal and 4 vertical arms. We then quantified the similarity between mazes by the cosine angle between pairs of vectors. A comparison with panels c and d indicates that the block-structure similarity inferred from the neural responses (f and h) is not due to maze geometry. **(d)** Average firing rate of two example neurons in Monkey F. Trials were sorted in 6 groups of 4. The 6 groups correspond to the 6 maze geometries numbered 1 to 6 (see panel b), and the 4 panels correspond to the 4 exit points. In each panel, the average firing rate is shown throughout the trial, from when the ball starts to drop until after the 3^rd^ flash with the flashes marked by three vertical lines (black: flash 1; dark gray: flash 2; light gray: flash 3). Strong similarities in response profiles are evident across mazes 1-3 (yellow background), and across mazes 4-6 (magenta background), but different across the two groups. Firing rates were computed from spike counts in 1-ms bins and smoothed with a truncated Gaussian kernel (σ = 100 ms). **(e)** Same as in panel (d) for two example neurons in Monkey N. The only difference is that the similarities between response dynamics form different groupings: similar across mazes 1-4 (yellow background), and across mazes 5-6 (magenta background), but different across the two groups. **(f)** Similarity matrix for response dynamics across maze geometries. For each neuron, we computed cross-validated pairwise correlations between activity profiles associated with the left-down condition that had identical arm lengths across all mazes (see Methods). We organized the pairwise correlations in a 6×6 similarity matrix, and averaged the values across all neurons (*n=1283*). Orange and Magenta square outlines are added for visual aid to dissociate between dynamical regimes. **(g)** Population neural trajectories projected on two Isomap dimensions for mazes 1 and 6 exemplifying the two clusters inferred from similarity matrix in (f). **(h)** Same as (f) for Monkey N (*n = 1570*). **(i)** Same as (g) for Monkey N.

For physiology experiments, we focused on a subset of six H-maze geometries (Fig. 2b) chosen to span the key features of the task space. Horizontal arm lengths were selected to produce three levels of left–right discriminability: ΔH=500 (*t_L_*=500 ms, *t_R_*=1000 ms), ΔH=250 (*t_L_*=500 ms, *t_R_*=750 ms), and ΔH=0 ms (*t_L_*=500 ms, *t_R_*=500 ms). The four vertical arms took values from the set {500, 700, 900, 1100 ms} and were arranged in two configurations. In the “paired” configuration, both vertical arms on one side were longer than the two vertical arms on the other side (*t_LU_*/*t_LD_*=900/1100 ms; *t_RU_*/*t_RD_*=700/500 ms), making vertical lengths informative about the left–right decision. In the “mixed” configuration, vertical arm lengths were interleaved across sides (*t_LU_*/*t_LD_*=700/1100 ms; *t_RU_*/*t_RD_*=900/500 ms), making them less readily helpful for left/right decisions. These manipulations were designed to create conditions under which different strategies would be advantageous, while avoiding distinct visual categories (Fig. 2c). Animals performed these mazes with high accuracy (Fig. S4). To ensure continued behavioral flexibility, a subset of trials (10%) included randomly generated arm lengths.

In principle, all six mazes could be solved using a single fixed strategy (Fig. 1i). However, we designed the geometries such that different mazes could elicit different strategies. For example, in mazes with large horizontal differences (ΔH = 500 ms), the left–right decision is relatively easy, making a hierarchical strategy effective by eliminating half of the options early. In contrast, in mazes with no horizontal difference (ΔH = 0 ms), the first interval is uninformative, and strategies that incorporate *t_2_* earlier such as postdictive strategies can improve performance. Alternatively, animals could ignore *t_1_* altogether and base their decision solely on the vertical arms. Thus, although a single strategy is sufficient in principle, the task structure creates conditions under which different strategies are advantageous.

Across 61 sessions, we recorded a total of 7,242 DMFC units (Monkey F: 3,429 units across 27 sessions; Monkey N: 3,813 units across 34 sessions). We first quantified the sensitivity of individual neurons to task variables using an ROC-based analysis (Fig. S5). Firing rates across a substantial fraction of neurons were modulated throughout the trial and could discriminate between the horizontal and the vertical arms (Table S1).

Having established a strong task-relevant modulation in DMFC, we carefully examined the response profile of individual neurons across the 24 task conditions (6 geometries x 4 exits). Task-modulated neurons had heterogeneous responses with complex and nonlinear dynamics. Some neurons showed transient responses aligned to specific task events, whereas others displayed sustained ramping activity that tracked observed or anticipated flashes (Fig. 2d,e).

Across the 24 conditions, response profiles exhibited a striking maze-dependent structure in both animals. In monkey F, two distinct groupings emerged: one associated with mazes 1–3 (Fig. 2d, yellow) and another with mazes 4–6 (Fig. 2d, magenta). In monkey N, a similar block structure was evident, with one grouping spanning mazes 1–4 and another spanning mazes 5–6 (Fig. 2e).

To quantify this organization, we compared neural responses across mazes using a similarity analysis. Direct comparison of response profiles was not possible because of differences in arm lengths and flash timing across mazes. However, from the outset, we designed the mazes with identical left–down (LD) arm lengths (*t_L_*=500 ms, *t_LD_*=1100 ms). We therefore focused our analysis on the response profiles of trials in which the ball took the LD path. We computed a similarity matrix by averaging the pairwise correlation of individual neurons’ responses across the 6 mazes (see Methods). The resulting similarity matrices revealed clear block structure in both animals (Fig. 2f,h), consistent with the animal-dependent groupings observed at the single-neuron level.

Importantly, this organization did not reflect visual similarity between maze geometries (Fig. 2c), indicating that the blocks reflect the structure of the underlying maze-dependent neural dynamics. Together, these results suggest that neural activity is organized into distinct geometry-dependent regimes.

To gain insight into the nature of these geometry-dependent regimes, we examined the structure of population dynamics using dimensionality reduction techniques, including principal component analysis (PCA; Fig. S6) and Isomap (Fig. 2g,i; Fig. S7). Across methods, neural trajectories exhibited distinct geometries corresponding to the two groups of mazes.

In one regime, trajectories diverged following the second flash, consistent with an early left–right commitment as expected under a hierarchical computation. In the other regime, trajectories remained largely overlapping after the second flash and instead diverged later in the trial, consistent with a sequential computation based on the time of the third flash. For example, in maze 1, neural trajectories showed an early bifurcation after the second flash, whereas in maze 6, trajectories were relatively insensitive to the second flash and diverged only after the third flash (Fig. 2g,i).

These observations indicate that the two regimes reflect distinct computational processes governing decision formation. The structure of these dynamics is consistent with hierarchical and sequential strategies, providing an initial link between geometry-dependent neural regimes and candidate solution strategies. Control analyses separating trials by choice confirmed that these patterns were not driven by averaging across heterogeneous trajectories (Fig. S8).

### Decoding computational algorithms

High-density Neuropixels recordings provide access to moment-to-moment fluctuations in neural population states ^84,85^. To characterize how these dynamical regimes unfold over time, we trained a decoder to predict left–right choices (marginalizing over up–down) from DMFC population activity in a 300 msec window after the third flash. The decoder achieved high accuracy on held-out trials and generalized to the 10% of trials with randomly sampled maze geometries, indicating that it captured robust, task-relevant signals (Fig. S9).

We next applied this decoder across the full trial to obtain a time-resolved estimate of the underlying left–right decision variable (DV). Specifically, at each time point, population activity was projected onto the decoder axis, yielding a continuous measure of the evolving decision state (Fig. 3). This approach provides a low-dimensional readout of population dynamics aligned to behaviorally relevant computations.

**Figure 3.**
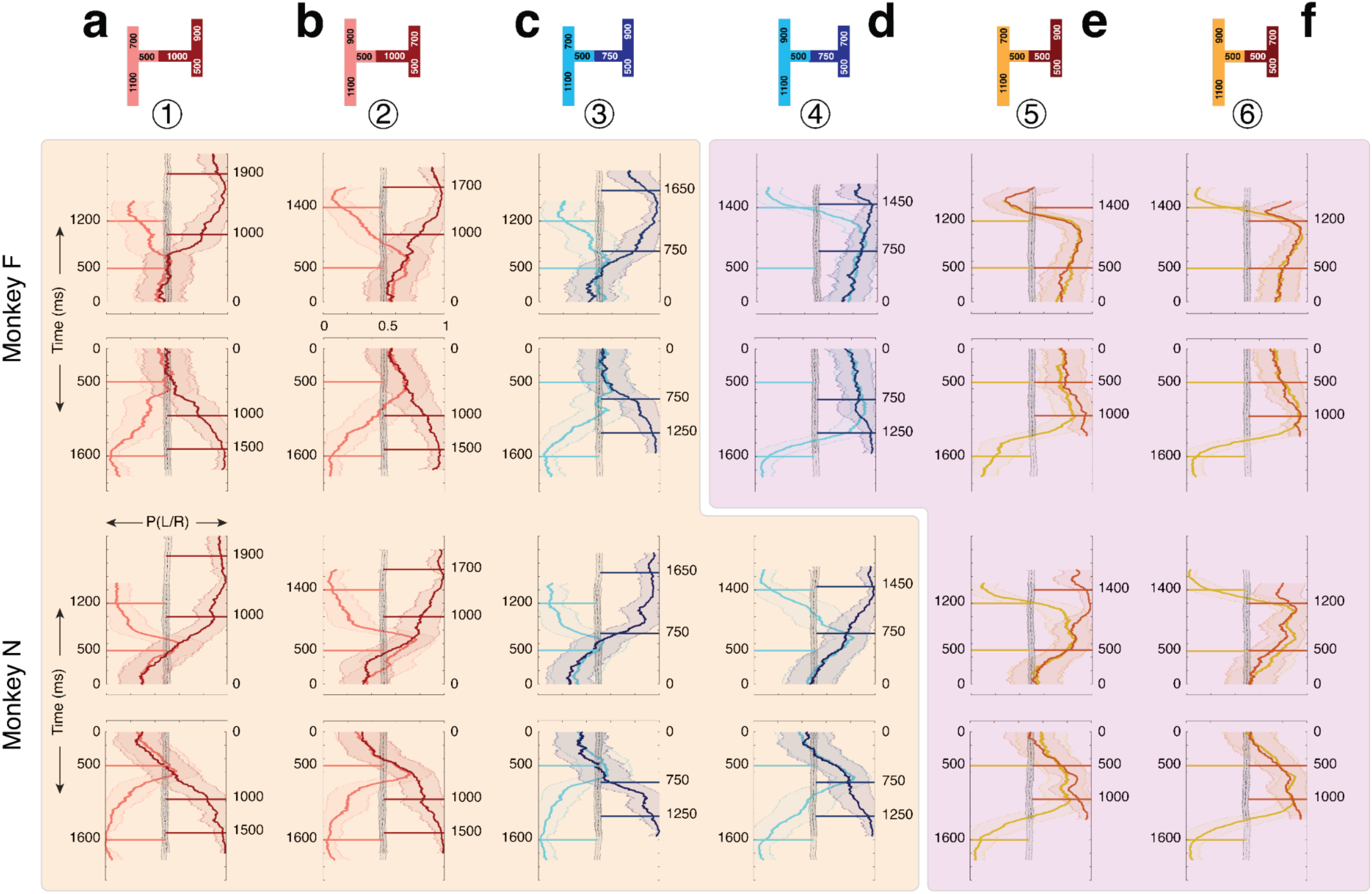
Left–right decision variable (DV) dynamics across maze conditions and animals. Results are shown separately for the two animals (monkey F: top; monkey N: bottom) across six maze geometries (columns a–f; mazes). Results are shown for the 10 highest-performing decoding sessions from each animal (see Methods). For each maze, DV trajectories for the four possible exits are shown in two panels (top panel: left-up and right-up exits, bottom panel: left-down and right-down). The DV (x-axis) is plotted in terms of probability of choosing right (left = 0, right = 1, chance = 0.5) as a function of time (y-axis) from the first flash until 300 ms after the third flash. Time increases upward/downward for top/bottom exits. In each panel, mean DV (solid lines) ± 0.5 standard deviations across trials (shaded areas) are color-matched to maze paths (top). Each panel also shows trial-shuffled mean DV (black) ± 0.5 standard deviations across trials (gray). Horizontal solid lines mark the timing of the second and third flashes for each exit. Background shadings separate mazes based on strategy (yellow: hierarchical, magenta: sequential).

We first examined DV dynamics on correct trials. Across mazes, the temporal evolution of the DV recapitulated the two geometry-dependent regimes identified above (Fig. 3a–f). In one regime, the DV diverged shortly after the second flash, consistent with early commitment to a left–right decision (Fig. 3a–c, top for monkey F; Fig. 3a–d, bottom for monkey N). In the other regime, the DV remained near baseline after the second flash and diverged later in the trial, indicating that the decisions were made based on the second interval (Fig. 3d–f, top for monkey F; Fig. 3e–f, bottom for monkey N).

### Continuous evolution of the decision state

Across conditions, the DV evolved smoothly throughout the trial, rather than changing only at the times of sensory flashes (Fig. 3a–f). This continuous evolution indicates that DMFC activity reflects a time-varying internal estimate of the decision state, consistent with ongoing integration of temporal information rather than discrete, event-triggered updates. Importantly, the smooth dynamics argue against a model in which decisions are computed elsewhere and only relayed to DMFC as discrete outcomes; instead, they suggest that DMFC reflects the ongoing formation of the decision ^86^.

### Predictive dynamics

The DV dynamics were predictive. For example, in monkey N, the DV initially favored the shorter horizontal arm and then shifted as time elapsed, crossing the decision boundary near the expected timing of the alternative (Fig. 3a–d, bottom). Shortly after the expected flash time, the DV reversed direction if the anticipated event did not occur, effectively ruling out one alternative before committing to the other. Similar, though weaker, patterns were observed in monkey F (Fig. 3a–c, top).

A related pattern was observed in mazes 4–6, where the early DV bias could not be explained by horizontal arm differences alone. In these mazes, the DV initially favored the right side despite identical or shorter horizontal arms on the left (e.g., Fig. 3d–f, Monkey F). This bias is consistent with anticipating the shortest vertical arm (500 ms), which was on the right. These results reveal that DV is governed by predictive dynamics.

### Inference by exclusion

DV dynamics also revealed sensitivity to the absence of expected events. In mazes 5 and 6, when flashes corresponding to one set of vertical arms failed to occur, the DV exhibited abrupt shifts favoring the alternative (Fig. 3e,f). This pattern indicates that animals used the absence of expected sensory events as evidence, effectively performing inference by exclusion. These dynamics extend beyond simple sequential evaluation and reflect a more flexible use of temporal information to guide decisions.

### Structured error patterns

We next examined DV dynamics on error trials (Fig. S10). Despite fewer error trials in some conditions (Fig. S4), the decoder still captured the overall decision dynamics. Critically, DV tracked the animal’s choice rather than the true stimulus: when animals made erroneous left–right decisions, the decoded state evolved toward the chosen—albeit incorrect—alternative. This indicates that the decoder captures the animal’s internal decision state rather than veridical task variables.

Across mazes, error dynamics exhibited two distinct patterns. In some cases, DV trajectories on error trials closely resembled those observed on correct trials, suggesting that the same underlying strategy was engaged and that errors arose from noise in internal timing estimates. In other cases, however, error dynamics differed qualitatively from the dominant pattern. For example, in monkey N, maze 4 showed hierarchical-like dynamics on correct trials but sequential-like dynamics on error trials, consistent with the use of a less adaptive strategy in that condition.

Together, these results suggest that errors arise from at least two sources: variability within a given strategy and trial-to-trial variation in strategy selection. This dissociation further supports the view that DMFC dynamics reflect the unfolding computational process underlying decisions and motivates the need for trial-by-trial inference of strategy.

### Unsupervised analysis of single trial trajectories

Our decoding analysis provided a time-resolved readout of decision dynamics. However, because decoding relies on projections onto a predefined axis, it does not capture the full structure of population activity. To characterize neural dynamics at the level of correct single trials, we used MARBLE ^87^, a nonlinear embedding approach that infers local flow fields from neural activity and represents them in a low-dimensional latent space, such that similar dynamical patterns are placed close together and dissimilar ones are separated (Fig. 4a–c).

**Figure 4.**
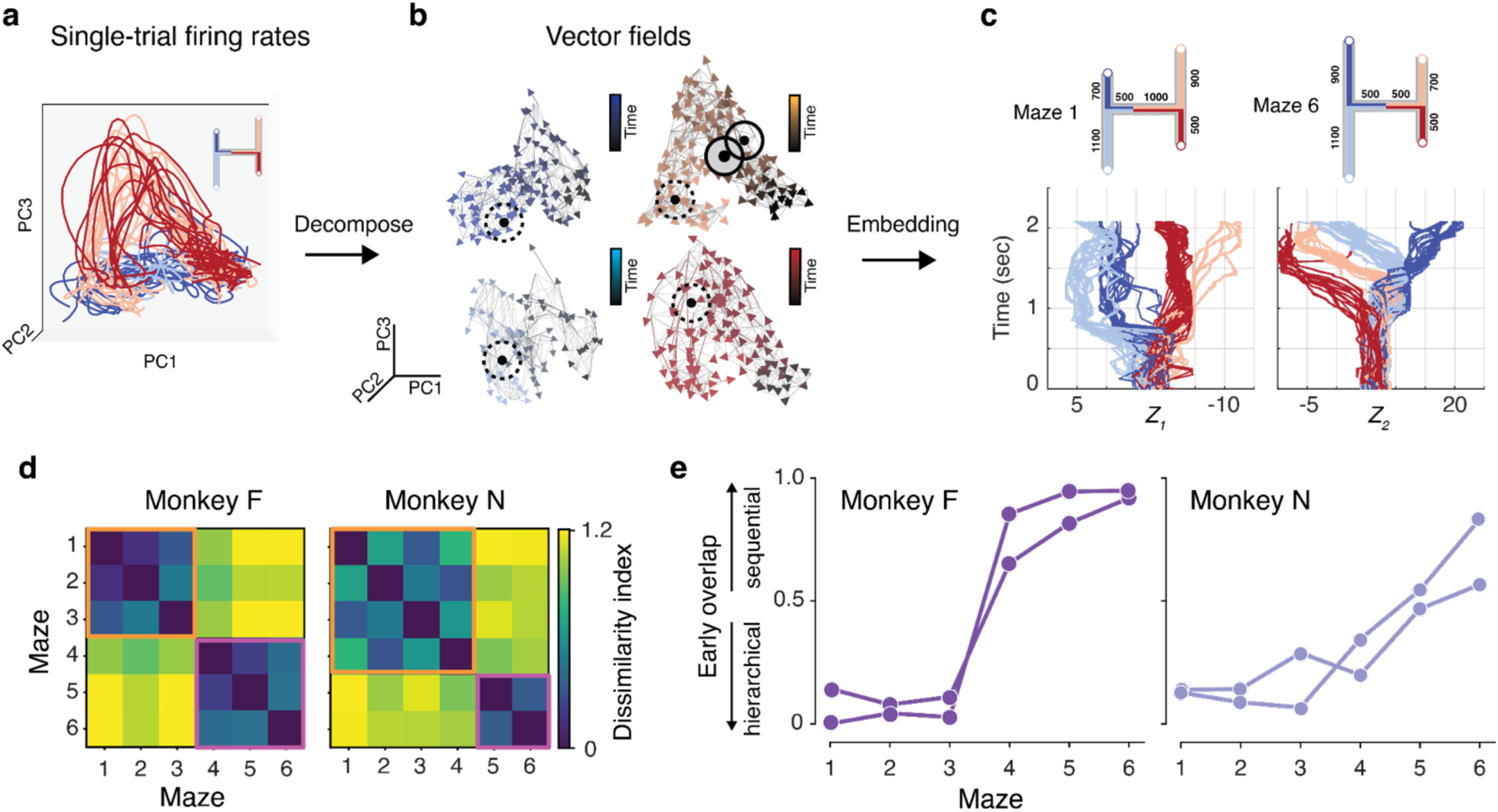
Unsupervised visualization and quantification of flow fields associated with correct single-trial neural population dynamics. **(a)** Single-trial spike times on correct trials across ball paths for monkey F and maze 1 (40 trials) visualized by three-dimensional PCA. **(b)** Flow fields inferred from MARBLE (subsampled for illustration). Arrows indicate direction of flow of neural states, shaded by trial time from black to the color of the respective ball path in a. The gray lines show a network indicating adjacency in neural space. Circles highlight examples of nearby (solid) and far apart (dashed) local flow fields (LFFs). **(c)** Embeddings of single-trial dynamics for mazes 1 and 6, visualized in one UMAP coordinate against time (maze 1: 40 trials, maze 6: 65 trials). **(d)** Heatmap showing the earth mover distance between the embeddings of the LD ball path for all maze pairs (monkey F: 292 correct trials, monkey N: 389 correct trials). Orange and Magenta square outlines are added for visual aid to dissociate between dynamical regimes. **(e)** Overlap between embeddings of the left (LU, LD) and right (RU, RD) ball paths within the time window of 300 ms after the second flash. We defined overlap based on the number of intersecting ε-balls centered at each latent state (see Methods). The results show average overlap across pairs of conditions (LU-RU, LU-RD, LD-RU, LD-RD). Small overlap indicates early bifurcation consistent with the hierarchical strategy. Large overlap indicates overlapping trajectories (no bifurcation) consistent with the sequential strategy. Results are shown for the two sessions from each animal with the highest session-quality metric (see Methods, monkey F: 292 (June 24) and 253 (June 8) correct trials, monkey N: 389 (November 3) and 345 (October 22) correct trials).

This approach revealed clear separation between the two dynamical regimes at the level of single trials. For example, embeddings for mazes 1 and 6 formed distinct trajectories with different temporal organization, consistent with the hierarchical and sequential patterns observed in both population averages and decoding analyses (Fig. 4c). To quantify this structure, we compared embeddings across mazes using the same dissimilarity metric applied to firing rates, focusing on the left–down (LD) path, which was identical across conditions. The resulting dissimilarity matrices exhibited a clear block structure in both animals (Fig. 4d), mirroring the previously identified organization (Fig. 2f-i). These results demonstrate that the two dynamical regimes are present at the level of single trials, consistent with the use of distinct strategies.

A key distinction between the two dynamical regimes is how neural trajectories behave around the time of the second flash. In one regime, trajectories diverge shortly after the second flash, whereas in the other they remain overlapping and diverge only later in the trial. We therefore reasoned that the degree of overlap between neural trajectories following the second flash could serve as a quantitative signature of the underlying computation.

To test this idea, we measured the overlap between neural embeddings associated with different trajectories within a 300 ms window after the second flash, separately for each maze (see Methods). Lower overlap reflects early bifurcation, consistent with the hierarchical regime, whereas greater overlap indicates that trajectories remain similar following the second flash, consistent with the sequential regime.

Across mazes, overlap increased systematically from maze 1 to 6 in both animals (Fig. 4e), indicating a gradual shift in the underlying dynamics from early to late divergence. These results provide a quantitative characterization of the transition between the two dynamical regimes and further support the interpretation that they reflect distinct computational processes.

### Behavioral validation of the maze-dependent mixed strategy

We next asked whether the geometry-dependent dynamical regimes identified in neural activity correspond to behaviorally relevant strategies. Specifically, we tested whether animals’ choices were better explained by a model in which strategy depends on maze geometry, as inferred from neural activity, or by models that assume a single fixed strategy across all trials.

To directly compare these alternatives, we focused on conditions in which the predictions of the fixed-strategy and neurally inferred mixed-strategy models diverged. For each maze, we assigned a strategy based on the average decoding results (Fig. 3), and constructed a mixed model that applied this strategy to generate behavioral predictions. We then compared this model to fixed hierarchical and fixed sequential models, restricting the analysis to trials in which the models made different predictions, thereby providing a stringent test of explanatory power.

Across both animals and both model comparisons, the neurally inferred mixed-strategy model consistently provided a better account of behavior, assigning lower negative log likelihoods than the two fixed-strategy models (Fig. 5a). This improvement was evident across individual maze conditions and in the overall averages for each animal, indicating that animals’ choices are more accurately captured by a model in which strategy varies with maze geometry.

**Figure 5.**
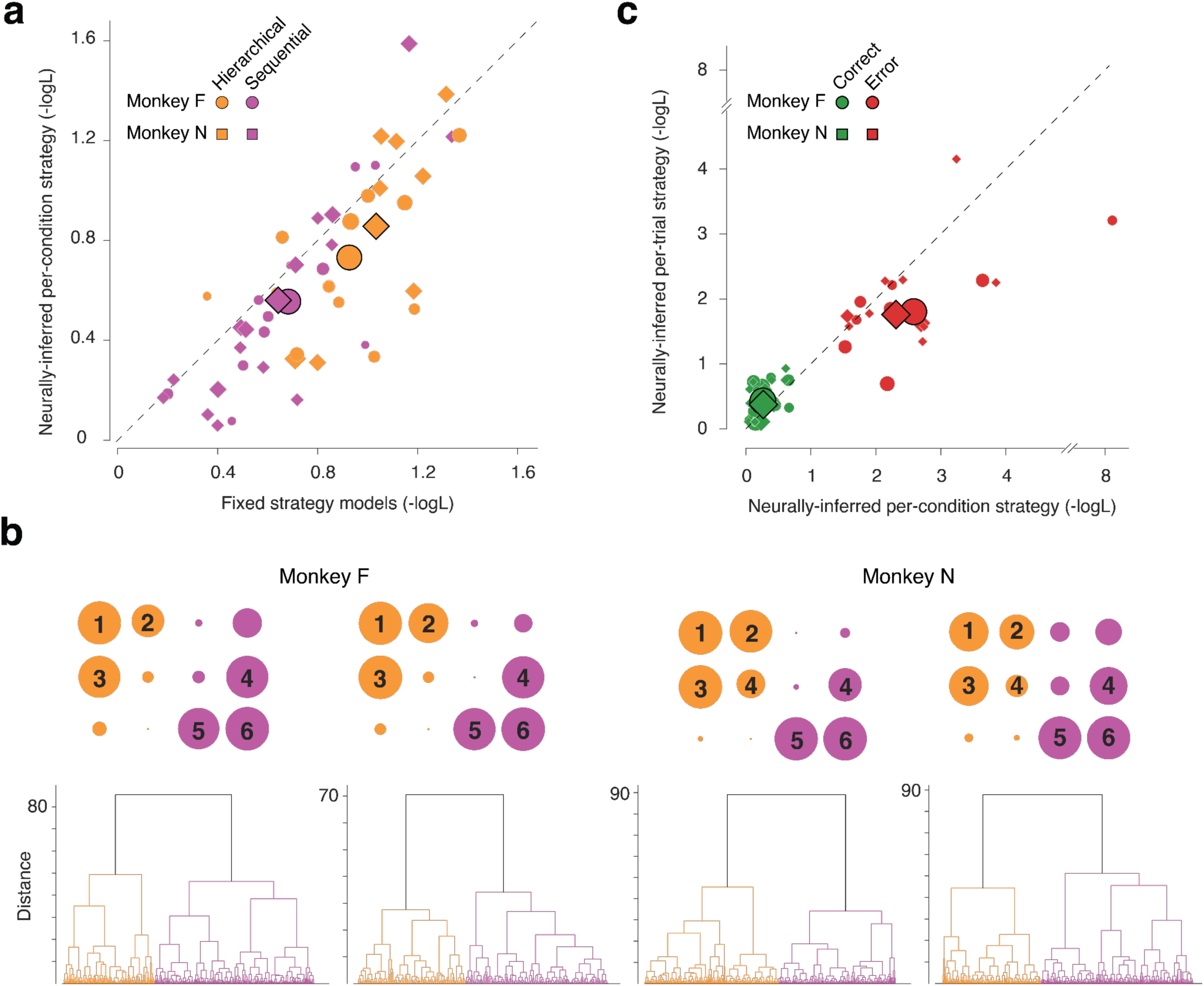
Linking initial neural states to strategy to behavior. (a) Behavioral predictions for the fixed strategy models versus a mixed strategy model in which maze-dependent strategies are inferred from decoding average neural data (Fig. 3). The mixed strategy model predicted behavior significantly better than either fixed strategy model (mixed versus sequential fixed model: left-tailed paired t-test, p = 3.468 × 10⁻^7^, n = 892; mixed versus hierarchical fixed model: p = 3.491 × 10⁻^14^, n = 750). Symbols show average per-path negative log-likelihood (Monkey F: circles, Monkey N: squares; Hierarchical: yellow, Sequential: magenta). Symbol size is proportional to the number of trials per path relative to the total number of trials for that monkey. Correct and error trials are pooled. Symbols with black outline indicate the overall mean values. The dashed line indicates unity. Results are shown for the two neurophysiology sessions with the highest data quality (see Methods). Only conditions in which the two models make different behavioral predictions are included. (b) Strategy-dependent organization of initial neural states. Bottom: Classification of initial neural states using an unsupervised hierarchical clustering algorithm for two sessions for each animal based on Euclidean distance (ordinate) in the state space (see Methods). Top: The contribution of trials from each maze geometry (enumerated 1-6) to the two top branches (top). The area of each circle is proportional to the number of trials in that maze geometry. Branches and leaf nodes are colored by assigned strategy. Results correspond to two sessions from Monkey F (June 24: 138 hierarchical, 243 sequential; June 8: 129 hierarchical, 171 sequential) and two from Monkey N (November 3: 270 hierarchical, 235 sequential; October 22: 183 hierarchical, 282 sequential). (c) Behavioral predictions of the condition-dependent mixed strategy model versus a model in which strategies are inferred on a trial-by-trial basis from the initial neural states. The per-trial strategy inferred from initial neural states predicted behavioral choices significantly better (left-tailed paired t-test: all violating trials, p = 0.0203, n = 226; error trials, p = 4.211 × 10⁻^7^, n = 62). Symbols show average per-path negative log-likelihood (Monkey F: circle, Monkey N: square; Correct trials: green, Error trials: red). Symbol size is proportional to the number of trials contributing to each per-path mean. For each monkey, marker sizes were scaled using path counts pooled across correct and error trials. Symbols with black outline indicate the overall mean values. The dashed line indicates unity. The difference between the two models was strongly driven by error trials. Results are shown for the two neurophysiology sessions with the highest data quality (see Methods). Only conditions in which the two models make different behavioral predictions are included.

### Early strategy selection is reflected in initial neural states

If the distinct dynamical regimes correspond to alternative strategies, then these strategies must be selected at the outset of each trial, based on the geometry of the maze. This predicts that neural activity prior to the onset of informative temporal cues should already reflect the upcoming computational mode.

To test this idea, we analyzed population activity during an early time window following maze presentation but before the ball reached the horizontal arm (see Methods). For each trial, we constructed population vectors from this initial period and performed hierarchical clustering across all trials. This analysis revealed a prominent binary split in the dendrogram for both animals (Fig. 5b), suggesting the presence of two dominant initial states.

Strikingly, the composition of these clusters closely matched the geometry-dependent groupings observed in the preceding analyses. Across sessions, clusters were dominated by distinct sets of mazes, with one cluster primarily containing mazes 1–3 and the other mazes 4–6 in monkey F, and mazes 1–4 versus 5–6 in monkey N (Fig. 5b, top). Thus, the initial neural state already reflects the structure of the maze and predicts which dynamical regime will unfold during the trial. Consistent with this, example dendrograms from individual sessions revealed a dominant binary split aligned with the inferred strategy identity (Fig. 5b, bottom).

This organization was also evident in the geometry of neural trajectories. Dimensionality reduction using PCA (Fig. S6) revealed that trajectories corresponding to different mazes occupied distinct regions of state space already at the time of the first flash, before any temporal evidence was available. A similar structure was observed using Isomap (Fig. S7), indicating that the separation of initial conditions is robust across methods.

Importantly, the clustering analysis also revealed variability at the level of individual trials. Although most trials from a given maze were assigned to a dominant cluster, a subset of trials was grouped with the alternative cluster, and this assignment was consistent with the predicted neural dynamics (Fig. S11). This trial-by-trial variability suggests that animals do not rigidly apply a single strategy to each maze, but instead occasionally switch between alternative strategies.

### Inferring strategy from single trial initial states

If these neurally inferred initial states reflect the strategy used on each trial, then they should provide a better account of behavior on trials in which they diverge from the maze-dependent mixed-strategy model. This possibility was suggested by the decoding analysis of error trials (Fig. S10), which indicated that some errors may arise from trial-to-trial variation in strategy selection. To test this, we focused on trials in which the strategy inferred from the initial neural state differed from the strategy assigned to that maze based on the average decoding results (Fig. 3). These trials provide a stringent test of whether single-trial neural signals capture the strategy used.

We compared the predictive power of two models: a mixed model in which strategy is assigned based on maze geometry, and a model in which strategy varies on a trial-by-trial basis according to the initial neural state. The neural model provided a better account of behavior overall (Fig. 5c). This advantage was driven primarily by error trials, where higher negative log likelihoods afforded greater sensitivity in distinguishing between the models. On correct trials, the mixed model performed comparably or marginally better.

These results show that trial-by-trial variations in behavior—particularly those that deviate from the dominant maze-dependent pattern—are captured by variability in neural initial conditions, indicating that strategy selection is not fixed at the level of maze geometry but varies across individual trials.

## Discussion

Early work on the neurobiology of decision-making has largely focused on relatively simple tasks in which behavior can be explained by a single, often near-optimal, strategy ^32,35,37,39,42,43,88–90^. More recent studies have extended this framework to context-dependent decisions, where the relevant context is either explicitly cued or learned through experience ^55,61,62,64,65,67,71,91–97^. In both cases, however, the structure of the problem and thus the relevant strategy is effectively prescribed by the experiment. More recent behavioral studies have begun to examine open-ended tasks in which tackle problems using alternative strategies ^82,98–101^. However, these studies remain agnostic about the underlying neural mechanisms. Here, we sought to elucidate the neural mechanisms that enable us to internally structure problems and flexibly deploy different strategies to solve them.

To tackle this question, we developed the H-maze task. This task is challenging and monkeys could solve it near-optimally. However, they grasped the logic of the task and were able to solve it at performance levels comparable to humans ^82^. Moreover, they were able to handle various parametric and structural generalizations, indicating that they could adapt their approach depending on the geometry of the maze. However, their choice patterns deviated from all fixed strategy and heuristic models we tested. It was only by examining neural activity that the underlying computational organization became apparent.

Large-scale recordings from the frontal cortex revealed two distinct dynamical regimes that corresponded to different modes of computation. These regimes were characterized by rich dynamical motifs, including dynamic elimination of upcoming options and inference by exclusion based on the absence of expected events. Such structure was not evident from behavior alone, but emerged clearly in the population dynamics. These results highlight the role of neural population activity as a window into the computational processes underlying behavior, and suggest that understanding higher cognition may require moving beyond behavioral observations ^102^, and require direct access to these dynamics.

A key finding of this study is that the initial state of the neural population predicts which dynamical regime will unfold during the trial. This indicates that the selection of a computational strategy occurs prior to the accumulation of task-relevant evidence. In other words, the animal appears to form a plan based on the geometry of the maze, and then execute that plan through the appropriate dynamical regime. The application of dynamical systems in neuroscience has already highlighted the importance of initial states as an important knob influencing neural computations supporting sensorimotor function ^103–106^. Our work demonstrates how the brain leverages initial states to select a specific computational strategy before tackling a problem. Notably, this form of deliberate strategy selection differs qualitatively from stochastic switching between decision policies, such as win–stay/lose–shift or motor biases ^107–109^. In H-maze, the initial state reflects the prospective selection of a rational strategy.

The comparison between maze-dependent and neurally inferred models further supports this interpretation. A model based on the dominant strategy for each maze accounted well for correct choices, whereas a model based on the initial neural state provided a better account of behavior on trials that deviated from this dominant pattern. In other words, although each maze is associated with a dominant strategy, the selection process is not deterministic: trial-by-trial variability in the initial neural state reflects variability in the selected strategy, which in turn predicts deviations in behavior.

While our results identify two dominant strategies—hierarchical and sequential—we do not yet understand why these particular strategies were selected, nor what determines their relative prevalence across animals. The two animals exhibited different partitions of maze geometries into these regimes, indicating that the mapping from task structure to strategy is not fixed. Notably, the animal that more frequently treated intermediate geometries (e.g., maze 4) as hierarchical also exhibited higher temporal acuity, suggesting that individual differences in timing precision may influence which strategies are advantageous. Therefore, strategy selection may reflect a form of metacognitive control that balances the expected costs and benefits of alternative computations under uncertainty ^18,22,110–112^. In the H-maze task, maze geometry modulates both the informativeness and timing of evidence, potentially favoring strategies that commit early when discriminability is high or defer commitment when it is low. Under this view, the observed strategies may emerge from a resource-rational tradeoff between speed, accuracy, and computational complexity, highlighting that adaptive problem solving involves not only selecting actions but also selecting flexibly among alternative computational procedures.

Our results may have broader implications for how the brain instantiates and selects among alternative algorithms. In our task, the same problem can be solved using different strategies, each corresponding to a distinct pattern of neural dynamics, and the animal must determine how to solve each instance by selecting among these alternatives. This process is reminiscent of how agents interact with objects by assessing what actions they afford. In Gibson’s formulation, affordances are the action possibilities that the environment offers to an agent ^113^. By analogy, the structure of a problem may afford a set of cognitive operations, from which the brain selects those most suitable for the task at hand. Decision strategies can thus be viewed as cognitive affordances: the problem affords a set of computational strategies, and the brain selects which one to apply. Consistent with this view, animals exhibited structured exploratory eye movements prior to fixation, selectively sampling different arms of the maze before task engagement, suggestive of an early assessment phase akin to hypothetical reasoning. After receiving feedback, animals also generated counterfactual eye movements toward unchosen alternatives, consistent with post hoc evaluation of outcomes. These pre- and post-decision behaviors point to a broader role for metacognitive control in which the brain evaluates, selects, and potentially updates strategies based on both prospective and retrospective information. More broadly, this suggests a unifying principle in which affordance-like mechanisms extend from the sensorimotor domain to the cognitive domain, supporting flexible and deliberate selection of alternative computational algorithms.

## Materials and Methods

## 1 Experimental subjects and recording setup

All procedures were approved by the Committee of Animal Care at the Massachusetts Institute of Technology and conformed to NIH guidelines. Experiments were conducted in two adult male rhesus macaques (Macaca mulatta; monkeys F and N; 7.2 and 10.2 kg; 9 and 6 years old). Animals were head-restrained and seated in a dark, sound-attenuated room. Visual stimuli were presented on a 23-inch LCD monitor (Acer H236HL; 60 Hz; 1920 × 1080 resolution) positioned in front of the animal. Eye position was recorded at 1 kHz using an infrared eye-tracking system (Eyelink 1000, SR Research). Eye calibration was performed at the start of each session using a 3 × 3 grid spanning the task-relevant region. Stimulus presentation and reward delivery were controlled using MWorks (http://mworks-project.org). Extracellular neural activity was recorded using Neuropixels probes (Neuro-Pixels probe 1.0, IMEC) through a bio-compatible cranial implant. All task events were synchronized precisely using an experimentally triggered photodiode signal. Analysis of behavioral and neural data was performed using custom MATLAB code (Mathworks, MA) and Python.

## 2 Behavioral task

Animals had to report the final location of a ball after it traversed through a two-dimensional maze. All mazes had a vertical entry corridor followed by multiple paths made of multiple 90° turns or bifurcations, each of which terminated in a potential final exit point. Arm width for all mazes was set to 1°. The ball diameter was set to 1°. The background was black, the maze arms were gray, and the ball was white.

Each trial began with the presentation of a maze with a randomly selected set of arm lengths (1.5 s), along with circles marking the possible exit points (white; diameter 1°). A fixation point (red; diameter 0.5°) was then presented, and animals were required to acquire fixation within 10 s or the trial was aborted. After a 0.7 s delay, the ball began moving through the maze at constant speed.

Identical visual flashes (white annuli centered at fixation; inner diameter 0.5°, outer diameter 1.25°, duration 80 ms) were presented each time the ball reached a bifurcation point and when it reached the final exit. These flashes therefore marked successive positions of the ball along its trajectory.

After a random delay period with a uniform hazard (minimum 300 ms, mean 500 ms), the fixation point disappeared. Animals were then required to make a saccade to the exit point corresponding to the final position of the ball. The trial was aborted if no eye movement was made within 1000 ms.

The selected target changed color to provide feedback (green if correct, purple if incorrect), and correct responses were rewarded with juice. Trials were aborted if fixation was broken while the fixation point was present. Feedback was displayed for 1 s. Trials were separated by an inter-trial interval of 1.5 s, with an additional 3 s timeout following fixation breaks.

### 2.1 H-maze

In most experiments, we used an H-shaped maze with an entry corridor of 7° that branched into two horizontal arms, each of which further branched into two vertical arms. We use the labels *L* (left), *R* (right), *LU* (left-up), *LD* (left-down), *RU* (right-up), and *RD* (right-down) to denote these arms. These labels are used as subscripts for arm lengths (*a_L_*, *a_R_*, *a_LU_*, *a_LD_*, *a_RU_*, *a_RD_*) as well as the corresponding traversal times (*t_L_*, *t_R_*, *t_LU_*, *t_LD_*, *t_RU_*, *t_RD_*).

In this configuration, the animal was presented with three flashes, one when the ball entered one of the horizontal arms, one when it entered one of the connected vertical arms, and one when it reached the corresponding exit. We will refer to the interval between the first and second flashes as the first sample interval, denoted by *ts_1_*, and the interval between the second and third flashes as the second sample interval, denoted by *ts_2_*.

#### 2.1.1 H-maze geometry: Behavioral sessions

Across behavioral experiments, the six arms of the H-maze each took one of seven possible lengths ranging from 4° to 10°. To ensure adequate coverage of task difficulty, arm lengths were constrained such that the absolute differences between paired arms (*|t_L_-t_R_|*, *|t_LU_-t_RD_|*, and *|t_RU_-t_RD_|*) formed a six-point discrete uniform distribution spanning 0° to 5°. To promote flexibility, ball speed was randomly set to 7 or 10 deg/s. Analyses were restricted to the 10 deg/s condition (Monkey F: 23 behavioral sessions, 15,457 trials; Monkey N: 24 sessions, 13,480 trials) since this was the only speed used during electrophysiology trials.

#### 2.1.2 H-maze geometry: Electrophysiology sessions

For electrophysiology experiments, we used a fixed ball speed (10 deg/s) and restricted the number of maze geometries to increase statistical power for neural data analysis. In 90% of trials, we used a fixed set of six H-maze geometries (Fig. 2b). Horizontal arm lengths were selected to produce three levels of left–right discriminability: ΔH=500 (*t_L_*=500 ms, *t_R_*=1000 ms), ΔH=250 (*t_L_*=500 ms, *t_R_*=750 ms), and ΔH=0 ms (*t_L_*=500 ms, *t_R_*=500 ms). Vertical arms took values from the set {500, 700, 900, 1100 ms} and were arranged in two configurations. In the paired configuration, both vertical arms on one side were longer than those on the other (*t_LU_*/*t_LD_*=900/1100 ms; *t_RU_*/*t_RD_*=700/500 ms). In the mixed configuration, vertical arm lengths were interleaved across sides (*t_LU_*/*t_LD_*=700/1100 ms; *t_RU_*/*t_RD_*=900/500 ms). In 10% of these trials, the ball was visible throughout its traversal to help animals maintain motivation. The visible trials were excluded from further analysis. The remaining 10% of trials consisted of randomly generated mazes sampled as in the behavioral sessions and were included to promote behavioral flexibility.

## 3 Modeling Behavior

To examine behavioral strategies, we compared each animal’s choice patterns in the H-maze task to predictions from a large set of models implementing different strategies.

The fundamental source of uncertainty in animals’ behavior is their noisy sense of time. Consistent with scalar variability of timing ^114^, we modeled timing noise as a sample from a zero-mean Gaussian whose standard deviation scales with elapsed time, with constant of proportionality, *w_m_*. Under this model, when the actual time between two flashes is *t_a_*, the conditional probability of the interval measured by the animal, denoted *t_m_*, is as follows:

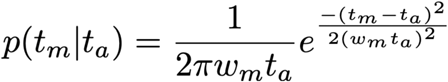

We estimated *w_m_* for each animal by analyzing the behavior in a simpler T-shaped maze. T-maze has an entry corridor that branches into two horizontal arms. In this configuration, animals see two flashes, one when the ball enters one of the horizontal arms and one when it reaches the corresponding exit. We denote the two arm lengths by their corresponding traversal times, *t_1_* and *t_2_* and the interval between the two flashes, *t_s_*.

To estimate *w_m_*, we formulated the decision policy using signal detection theory^115^ with two additional parameters, a criterion, *t_c_*, to account for each animal’s bias, and a lapse rate parameter, Γ. Without loss of generality, we use the convention that *t_1_* is the shorter of the two arms (i.e., *t_1_* ≤*t_2_*). Accordingly, the psychometric function can be written as follows:

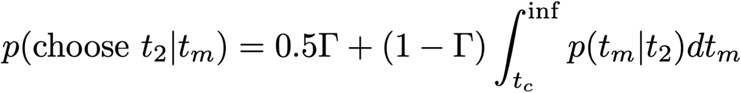

We used maximum likelihood to find the best *w_m_* (as well as the other two parameters) by fitting the psychometric function to each animal’s choices using a binomial distribution. We obtained robust parameter estimates by averaging across 100 bootstraps, each using 80% of trials (Monkey F: *w_m_* = 0.165, *t_c_* = 0, Γ = 0.0092, n = 708 trials; Monkey N: *w_m_* = 0.153, *t_c_* = 0, Γ = 0.0154; n = 517 trials). For plotting, we generated a smooth psychometric function using the mean bootstrap parameter estimates evaluated on a dense grid of arm-length combinations defining 100 evenly spaced values for each arm length parameter across the full range observed in the data. We then used *w_m_* to construct predictive behavioral models for each animal for the H-maze.

### 3.1 4AFC Models

#### 3.1.1 Optimal model

This model chooses the most likely exit by comparing *p(tm_1_,tm_2_|t_L_,t_LD_)*, *p(tm_1_,tm_2_|t_L_,t_LU_)*, *p(tm_1_,tm_2_|t_R_,t_RU_)*, and *p(tm_1_,tm_2_|t_R_,t_RD_)* where *tm_1_* and *tm_2_* are the two noisy measured intervals. We assumed that the two measurements are conditionally independent and computed the joint probabilities by the product of the marginals; e.g., *p(tm_1_,tm_2_|t_L_,t_LD_) = p(tm_1_|t_L_)p(tm_2_|t_LD_)*. Note that this model has no additional parameters other than *w_m_* that was estimated from the T-maze experiment.

#### 3.1.2 Optimal model with lapses

This model is the same as the optimal model except that it allows lapses at a rate of Γ when the model chooses randomly between the four exits (25% chance performance). We estimated Γ using maximum likelihood estimation.

#### 3.1.3 Total time

This model ignores the second flash and decides based on the total time between the first and third flash by comparing *p(tm_1_ + tm_2_|t_L_ + t_LD_)*, *p(tm_1_ + t m_2_|t_L_ + t_LU_)*, *p(tm_1_ + tm_2_|t_R_ + t_RU_)*, and *p(tm_1_ + tm_2_|t_R_ + t_RD_)*. The conditional probability for *tm_1_ + tm_2_* is the convolution of the individual conditional distributions centered on the sum of the respective means.

### 3.2 2×2AFC Models

#### 3.2.1 Hierarchical Model

This model compares *p(tm_1_|t_L_)* and *p(tm_1_|t_R_)* to make a left-right decision. Subsequently, it compares the conditional probability for the corresponding vertical arms: *p(tm_2_|t_LD_)* versus *p(tm_2_|t_LU_)* for left and *p(tm_2_|t_RD_)* versus *p(tm_2_|t_RU_)* for right. This model also has no additional parameter except *w_m_*.

#### 3.2.2 Postdictive Model

This model is similar to the hierarchical model except that it makes its initial left-right decision postdictively by incorporating additional information about the *tm_2_* and the vertical arms. Specifically, the model compares *p(tm_1_,tm_2_|t_L_,t_LD_)*+*p(tm_1_,tm_2_|t_L_,t_LU_)* to *p(tm_1_,tm_2_|t_R_,t_RU_)*+*p(tm_1_,tm_2_|t_R_,t_RD_)* to make a left-right decision. Subsequently, it compares the conditional probabilities for the corresponding vertical arms. Similar to the joint and hierarchical model, this model has no additional parameter.

#### 3.2.3 Hierarchical + Revision Model

This model is similar to the hierarchical model but can revise its initial left-right decision if its confidence in that choice is below a certain threshold. We constructed four variants of this model by considering different formulations of confidence (Table 1). The threshold parameter, *θ*, was fitted for each model variant independently. The results in Fig. 1i are based on the best-fitting hierarchical + revision model.

**Table 1.** Hierarchical + Revision Models. First column: Abbreviated names. Middle column: quantity used as confidence. Right column: Description of the confidence measure where chosen side refers to the initial left-right decision prior to the revision.

| Model Name | Confidence measure | Description |
| --- | --- | --- |
| $X_{UD}$ | $p(U \text{ or } D\vert tm_2)$ | Probability of either up or down |
| $X_{max}$ | $\max(p(U\vert tm_2) \& p(D\vert tm_2))$ | The larger of the two probabilities for up and down |
| $X_{LR/UD}$ | $p(L/R\vert tm_1) * p(U \text{ or } D\vert tm_2)$ | Same as $X_{UD}$ scaled by probability of chosen side |
| $X_{LR/max}$ | $p(L/R\vert tm_1) * \max(p(U\vert tm_2) \& p(D\vert tm_2))$ | Same as $X_{max}$ scaled by probability of chosen side |

An important consideration for this model is that revisions must incur a performance cost. If revisions were cost-free and lossless, the model could set the threshold to zero (“always revise”), effectively evaluating all options and becoming indistinguishable from the optimal model. To address this, we assumed that maintaining information about *tm_1_* and *tm_2_* in working memory during revision is lossy. Specifically, we assumed that *tm_1_* and *tm_2_* become noisier after each revision. We modeled the noise by a Gaussian distribution with mean *β* and standard deviation that is proportional to the measured interval (*tm_1_* or *tm_2_*) with a fixed constant of proportionality *α* (in accordance with scalar variability of time ^114^). The parameters *α* and *β* were fitted using maximum likelihood estimation.

### 3.3 Other heuristic models

We also considered two heuristic models, one that only relied on the first time interval between flash 1 and 2, and another that only relied on the second time interval between flash 2 and 3 (Table 2).

**Table 2.** Other heuristic models. Firm column: Model name. Second column: Decision policy.

| Model name | Decision policy |
| --- | --- |
| First-only | Compare $p(tm_1\vert t_L)$ and $p(tm_1\vert t_R)$ to make a left-right decision. Choose up-down randomly |
| Second-only | Choose exit by comparing $p(tm_2\vert t_{LU})$, $p(tm_2\vert t_{LD})$, $p(tm_2\vert t_{RU})$, and $p(tm_2\vert t_{RD})$ |

### 3.4 Model fitting

Many of the models did not have any additional parameters; their only parameter was *w_m_*, which we estimated from the T-maze experiments. Other models had additional parameters that were fit to behavior using maximum likelihood. Parameters were estimated using 70% of randomly selected trials for training, and model performance was evaluated on the remaining 30% held-out trials. This procedure was repeated across five random seeds, and model predictions for each run were generated using the corresponding maximum-likelihood parameter estimates and evaluated on the matched held-out trials.

We estimated choice probabilities using 1000 stochastic Monte Carlo simulations per trial and parameter setting. Probability for each choice was estimated as the fraction of simulations yielding that choice, with values floored at machine precision to avoid undefined likelihoods. These probabilities were then used to compute the multinomial negative log-likelihood of the animal’s observed choices, summed across trials under an independence assumption.

### 3.5 Comparing model to animals

We compared models and animals using a log performance ratio (Fig. 1j). Because our experiment involved an extremely large number of unique trial conditions (a total of 7^6^ unique conditions), we first grouped trials based on the absolute horizontal arm time difference (|*t_L_*-*t_R_*|) and the vertical arm time difference associated with the correct horizontal side (|*t_LU_*-*t_LD_*| or |*t_RU_*-*t_RD_*|), and then quantified the discrepancy between monkey and model behavior independently for each group (Fig. 1j). Model performance was estimated by averaging across five runs using the best-fitted parameters on training data and evaluated on held-out data. We assessed overall goodness of fit using the coefficient of determination (R^2^) between the monkey and model performances across the groups.

To identify conditions in which the model deviated significantly from animals, we compared the monkey performance in each condition to the distribution of model performance across runs using a two-sided one-sample t-test. To illustrate model failure (Fig. 1k), we plotted model and animal choice probabilities (*p_LU_*, *p_LD_*, *p_RU_*, and *p_RD_*) for the condition where the model and animals were most dissimilar. We quantified this dissimilarity using the Euclidean distance in a four-dimensional space spanned by *p_LU_*, *p_LD_*, *p_RU_*, and *p_RD_*.

## 4 Neurophysiology

We recorded extracellularly neural activity through a bio-compatible cranial implant targeting the dorsomedial frontal cortex (DMFC), comprising supplementary eye field (SEF), presupplementary motor area (Pre-SMA), and dorsal portion of the supplementary motor area (SMA) excluding the medial bank. Regions of interest were selected by cross-referencing structural scans with stereotaxic coordinates based on prior studies ^67,116–120^. For monkey F and N, we conducted 27 and 34 recording sessions yielding a total of 3429 and 3813 units, respectively.

Recorded signals were acquired using SpikeGLX, amplified, bandpass filtered, and sampled at 30 kHz. Spikes from single- and multi-units were sorted offline using Kilosort (v2.5). Following sorting, we applied a custom waveform-based curation procedure. For each cluster, channels were ranked by waveform amplitude, defined as the difference between the maximum and minimum of the mean waveform on each channel.

Clusters were labeled as noise if they met any of the following criteria: excessive sparsity across channels (top-to-second channel amplitude ratio > 3), spatial multimodality (top two channels separated by >10 channels), low amplitude (peak-to-trough < 16), excessive spatial spread (>30 channels exceeding 60% of the mean amplitude of the top three channels), or strong cross-channel waveform-shape dissimilarity (dissimilarity score > 0.4).

Waveform-shape dissimilarity was computed after mean subtraction and amplitude normalization, allowing circular temporal shifts to account for small timing offsets. For Neuropixels recordings, flagged clusters were relabeled with the corresponding exclusion reason and excluded from downstream analyses.

Units with fewer than 250 stable trials or a mean firing rate below 0.25 spikes/s were excluded. Spike counts were computed in 1-ms bins and smoothed with a truncated causal Gaussian kernel (σ = 100 ms) to characterize single-neuron response profiles.

### 4.1 Sensitivity analysis

We used a receiver operating characteristic (ROC) to quantify the degree to which each neuron could discriminate between pairs of horizontal and vertical arms. For the horizontal arms, we measured the area under ROC (AUC) using firing rate in 100 ms windows prior to the second flash comparing leftward and rightward trials (excluding mazes 5 and 6 in which the horizontal arm lengths were equal). For the vertical arms, we measured AUC based on firing rate in the 100 ms window preceding the third flash comparing upward and downward trials separately for trials in which the ball initially moved left or right. Firing rates were Z-scored independently for each neuron across all valid trials and time points within a session.

To avoid bias from class imbalance, trial counts were equalized by randomly subsampling the larger class to match the smaller class. Statistical significance was assessed using 100 label shuffles. Because our goal was to assess sensitivity irrespective of preferred direction, AUC values were treated as unsigned. Neurons were classified as significant if their observed AUC exceeded the 95th percentile of the shuffled distribution. Only neurons with at least two trials per condition across all 24 path conditions were included in the analysis.

### 4.2 Similarity analysis

To assess the similarity of single-neuron response profiles across maze conditions, we performed a split-half correlation analysis on trial-averaged firing-rate time courses from fixation onset to 300 ms after the third flash. Analyses were restricted to the correct left-down trials, which featured identical flash timings across all six mazes. For each neuron, trials from each maze were randomly divided into two halves over 10 iterations. Trial-averaged firing-rate time courses were computed separately for each half, and pairwise Pearson correlations were calculated between response profiles from one half of one maze and the other half of another maze. Correlation matrices were first averaged across split-half iterations for each neuron and then across neurons to obtain the final maze-by-maze similarity matrix. Analyses were restricted to neurons with at least 10 valid trials per maze and a mean firing rate above 1 spike/s in at least one maze. To ensure comparable trial counts, trials were randomly subsampled so that each maze contributed an equal number of trials, matched to the minimum across mazes.

### 4.3 Visualizing population activity

To visualize how neural population state evolved across task conditions, we analyzed trial-averaged neural trajectories aligned to key task events. For each analysis, time windows were selected to capture the relevant task epoch. Prior to dimensionality reduction, each neuron’s activity was mean-centered across all samples included in that analysis.

#### 4.3.1 Linear dimensionality reduction (PCA)

We first applied principal component analysis (PCA) to obtain a low-dimensional linear representation of population activity. For each analysis, population matrices were constructed by concatenating trial-averaged neural activity across conditions and time, with neurons treated as features and condition–time points as samples. Firing rates were Z-scored independently for each neuron across all valid trials and time points within a session.

For the initial visualization, we grouped trials based on path condition and applied PCA to neural activity from flash 1 to 300 ms after the third flash.

We also used PCA to examine the emergence and timing of early trajectory bifurcations. Analyses were restricted to trials in which the animal made the correct left–right decision, so that the trajectories reflected canonical path-related structure. For each path condition (LU, LD, RU, and RD), firing rates were averaged across these trials. PCA was applied to neural activity from flash 1 to 1,250 ms after flash 1, a fixed interval chosen to encompass the duration of the shortest path across mazes. Neurons with missing values (e.g., due to absence of trials for a given condition–choice combination) were excluded.

#### 4.3.2 Nonlinear embedding (Isomap)

To capture potential nonlinear structure in population activity, we applied Isomap to PCA-reduced responses. Firing rates were Z-scored independently for each neuron across all valid trials and time points within a session.

To visualize the topology of population dynamics for each maze, we applied Isomap separately to each maze (four possible ball paths) using the top 10 principal components, 150 nearest neighbors, and a two-dimensional embedding.

To visualize the shared structure, we applied Isomap to the pooled dataset using the top 5 principal components and 200 nearest neighbors.

### 4.4 Decision variable dynamics

To obtain a time-resolved decision variable (DV), we trained a decoder on endpoint population activity and projected neural activity throughout the trial onto this axis. We trained supervised logistic regression decoders of the animal’s final left–right choice based on average firing rates within a 300 ms window after the third flash.

Firing rates were Z-scored independently for each neuron across all trials within a session. Decoders were trained using MATLAB’s `fitclinear` function with a logistic learner, stochastic gradient descent optimization, ridge regularization, and a uniform class prior. The model minimized the logistic loss:

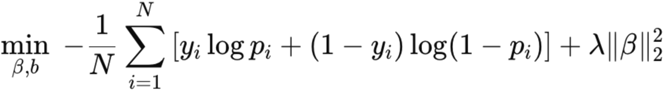

where *y_i_* denotes the behavioral choice label, *p_i_* is the logistic function, and λ is the regularization parameter. λ was selected via 10-fold cross-validation from a logarithmically spaced range (10^-6^ to 10^-0.5^), balancing classification accuracy and coefficient magnitude. The logistic function (*p_i_*) was defined as:

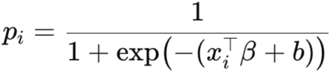

where *x_i_* is the population firing-rate vector, β the fitted weight vector, and *b* a scalar bias term.

We trained session-specific decoders on endpoint neural activity using 200 repeated 50% holdout iterations, balancing classes within each training set by subsampling. The decoder trained on endpoint activity was then applied to held-out trials across the full time course of the trial (from the first flash to 300 ms after the third flash) to generate a time-resolved decision variable (DV).To assess generalization, the same decoders were also evaluated, without retraining, on the 10% of trials in each session in which mazes with randomly sampled arm lengths were presented. These random-geometry trials were excluded from both training and held-out testing and were treated as an out-of-distribution validation set.

At each time point, population activity was projected onto the decoder axis to obtain a continuous estimate of the evolving decision state. Final decoding results were computed by averaging across iterations using held-out trials only.

### 4.5 Unsupervised representation learning

We used MARBLE, a geometric deep learning framework ^87^, to elucidate the distinct dynamics associated with different behavioral strategies from single-trial data.

First, we used Z-scored neural trajectories from correct trials, denoted by the vector *r*(*t*), to construct a difference vector field by taking temporal differences, *x*(*t*) = *r*(*t* + 1) − *r*(*t*). Next, we used the vector field to construct local flow fields (LFFs) at each neural state. To minimize biased sampling, we applied furthest point sampling ^121^ to generate an approximate uniform sampling of neural states. An average inter-state spacing of ∼3% of the manifold diameter yielded good coverage without excessive data exclusion. This procedure was repeated with three independent subsamplings. We then constructed a proximity graph over the subsampled states by connecting nearby states based on Euclidean distance. To ensure that the graph was insensitive to local sampling density, distances were normalized by the distance of each state to its *k-*th nearest neighbor ^122^. Specifically, states *r*_i_ and *r*_j_ were connected if:

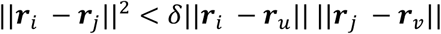

where ||. || is Euclidean distance, and *u* and *v* are the *k*th nearest neighbor of *i* and *j*, respectively.

Using this graph, we defined LFFs at each neural state as the collection of difference vectors that were within two hops of that state. We then used a geometric neural network to learn the coefficients of a Taylor series expansion (up to the second order) needed to approximate a latent vector *z_i_* that represents LFFs across the entire state space.

The network was trained via unsupervised contrastive learning, minimizing the loss function

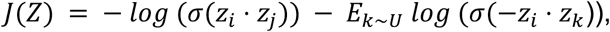

where σ(*x*) = (1 + *e*^-x^)^-1^ is the sigmoid function, *i*, *j* are neighboring nodes over the connectivity graph and *E*_k∼U_ is the expectation taken over nodes drawn uniformly from all nodes. This loss function ensures that similar LFFs remain close in latent space, thereby preserving similarities. For training we split the dataset into 80% training points, 10% validation points and 10% test points. We used the following hyperparameters to train MARBLE: *k=20,* δ=1.4, epochs=100, batch size=32, architecture: two-layer multilayer perceptron with 50 hidden units and output dimension 5. We found that these hyperparameters yielded well-structured latent representations, as indicated by a minimal gap between training and validation losses.

#### 4.5.1 Trajectory overlap metric

To quantify the extent to which neural dynamics reflected hierarchical versus sequential computations, we defined a continuous metric based on the overlap of left and right trajectories in the MARBLE latent space following the second flash. The key intuition is that hierarchical strategies produce an early bifurcation between left and right trajectories, whereas sequential strategies maintain overlapping trajectories until later in the trial.

We operationalized this idea by measuring the degree of overlap between latent states associated with left and right paths shortly after the second flash. Two latent states, *z*_i_ and *z*_j_ were considered overlapping if their distance was less than ε. The threshold ε was defined as the average distance between nearest-neighbor neural states prior to the second flash (< 500 ms), providing a data-driven estimate of baseline variability before trajectory divergence.

The overlap score was defined as the fraction of overlapping left–right neural state pairs:

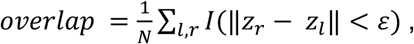

where *z*_l_and *z*_r_denote left and right neural states, respectively, *I* is the indicator function, returning 1 if the argument is true and 0 otherwise, and *N* is the total number of evaluated state pairs for left and right trajectories. Lower overlap indicates early separation of trajectories, consistent with hierarchical dynamics, whereas higher overlap reflects delayed divergence, consistent with sequential dynamics. Overlap was computed using latent states from the interval between the second flash and 300 ms afterward, and results were averaged across five independently trained MARBLE models.

### 4.6 Initial neural states

We used hierarchical clustering of single-trial population activity to test whether initial neural states formed maze-dependent clusters consistent with the ensuing dynamics. We focused on a time window after maze presentation but before the ball entered the horizontal segment.

Initial neural states were denoised using PCA. For each trial, firing rates were averaged over a 500 ms window following the first flash. Trials were pooled across path conditions and concatenated across mazes to form a trial-by-neuron matrix, which was z-scored across trials for each neuron and projected onto the first three principal components. Hierarchical clustering was then performed on this low-dimensional representation using Euclidean distance and Ward linkage.

The dendrogram was cut at the first split to define two clusters for downstream analyses. We quantified the distribution of trials from each maze across clusters and estimated chance levels by shuffling maze labels across trials 100 times and recomputing cluster assignments.

### 4.7 Data selection/exclusion criteria

Analyses relying on populations of simultaneously recorded neurons (Fig. 3-5) were restricted to the block of contiguous trials with stable recordings that retained as much of the session as possible. Specifically, for each candidate block, we computed the fraction of neurons present across all trials (ƒ_neurons_) and the fraction of trials retained (ƒ_trials_), and selected the block that maximized a retention score defined as ƒ_neurons_ + ƒ_trials_ − |ƒ_neurons_ − ƒ_trials_|). This score favors blocks that retain both many neurons and many trials while penalizing imbalances between the two.

For decoding results (Fig. 3), we focused on the 10 sessions from each animal with the highest end-point decoding performance. For unsupervised learning (Fig. 4) and initial condition results (Fig. 5), we focused on the top two sessions where the best neural data for mazes 1 and 6 clearly discriminated between the hierarchical and sequential strategies. To identify candidate sessions, neural activity was projected onto the first three principal components, and trials were classified using a linear support vector machine with stratified 5-fold cross-validation. We use ROC analysis to quantify the discriminability between the two strategies.

## 5 Behavioral models based in neurally inferred strategies

### 5.1 Fixed versus maze-dependent strategy

To compare fixed versus maze-dependent accounts of strategy, we evaluated the likelihood of each model in explaining the animal’s trial-by-trial choices. For each observed trial, model predictions were estimated by Monte Carlo simulation (10,000 samples) using the observed arm lengths and the monkey-specific timing variability parameter (*w_m_*), yielding a probability distribution over the four possible exits. Trial-wise negative log-likelihoods were computed from the probability assigned to the animal’s chosen exit, with probabilities floored at machine precision to avoid undefined values.

To ensure that comparisons were informative, analyses were restricted to trials in which the hierarchical and sequential strategies made different predictions. For the maze-dependent model, strategy was assigned at the level of maze geometry based on the dominant decoding pattern for each animal: for monkey F, mazes 1-3 were assigned to the hierarchical set and mazes 4-6 to the sequential set; for monkey N, mazes 1–4 were assigned to the hierarchical set and mazes 5 and 6 to the sequential set.

In contrast, fixed strategy models applied a single strategy across all mazes. Trial-wise likelihoods were pooled across sessions separately for each monkey and averaged within each of the 24 path conditions. Trials for which the model assigned zero probability to the observed choice were excluded from summary analyses. Statistical comparisons used one-tailed paired t-tests to test whether the maze-dependent model produced lower negative log-likelihood than the fixed model.

### 5.2 Maze-dependent versus trial-dependent strategy

We next compared a maze-dependent strategy model with a trial-dependent model in which strategy was inferred from the initial neural state on each trial. In the maze-dependent model, strategy was assigned at the level of maze geometry as described above. In the trial-dependent model, strategy labels were derived from cluster memberships at the first split of the single-trial dendrogram (see above, Initial neural states).

For each session, the two clusters were mapped onto hierarchical and sequential strategies based on their maze composition: the cluster containing more trials from maze 1 was labeled hierarchical, whereas the cluster containing more trials from maze 6 was labeled sequential. These two mazes lie at opposite ends of the stimulus space and provide the strongest separation between the two strategies. All trials within each cluster inherited the corresponding strategy label, yielding a trial-wise strategy assignment.

Analyses were restricted to violating trials, defined as trials in which the trial-dependent strategy differed from the maze-level assignment. Correct and error trials were analyzed separately. Trial-wise likelihoods were pooled across sessions within each monkey and summarized by averaging within each of the 24 path conditions. Trials for which the model assigned zero probability to the observed choice were excluded from summary analyses. Statistical comparisons used one-tailed paired t-tests to test whether the trial-dependent model produced lower negative log-likelihood than the maze-dependent model.

## Author contributions

M.R. and M.J. conceived the study. M.R. collected the data and performed all the analyses. A.G. and M.R. wrote the MARBLE training and evaluation code. M.R., A.G. and M.J. wrote the manuscript. M.J. supervised the project.

## Competing interest

The authors declare no competing interest.

## Data availability

The data used to generate the associated figures will be made available on a public repository after peer-review publication.

## Code availability

The code used to generate the associated figures will be made available on a public repository after peer-review publication.

## Acknowledgments

M.R. is supported by ICoN and Pillar/MIT Fellowship. A.G. acknowledges support from an ERC Starting Grant (NEURO-FUSE, Project DOI: 10.3030/101163046). M.J. is supported by the Simons Foundation. HHMI, and the McGovern Institute.

## Supplementary Material

**Supplementary Video 1**. Example H-maze trials from monkey F (correct feedback: green; incorrect feedback: purple).

## Supplementary Figures

**Supplementary Figure 1.**
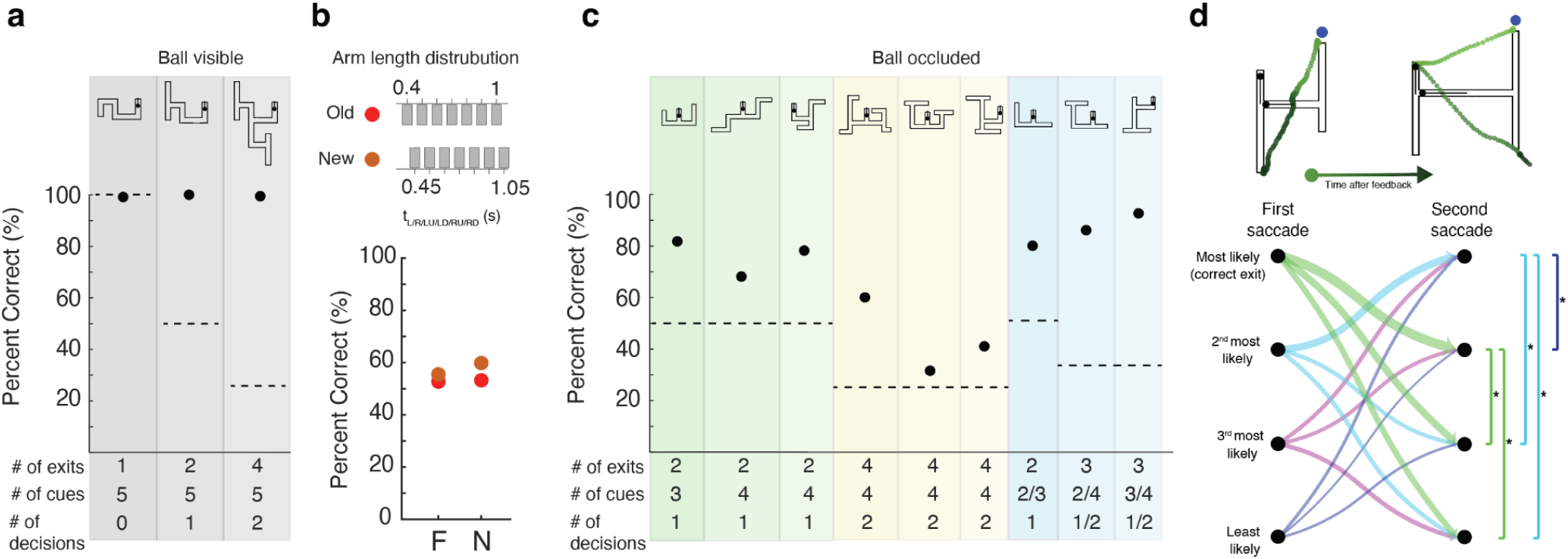
Generalization performance. **(a)** Mean performances averaged across monkeys (black dot) during generalization experiments with new maze geometries (shown on top) for when the ball was visible throughout its movement. The number of exits, cues (visual flashes), and decisions for each maze are shown below the abscissa. Dashed lines correspond to chance performance. **(b)** Mean performance of monkeys F and N for H-mazes for arm lengths drawn from the training distribution (top, red) as well as a new test distribution (top, orange). **(c)** Same as panel (a) for generalization experiments when the ball was occluded. **(d)** Top: Two example trials showing post-feedback eye movements relative to the maze (Monkey F, left; Monkey N, right). Animals made additional saccades after the first saccade (blue circle) as evident from eye position over time color-coded from light to dark green. In both examples, the top-left is the correct response. Left: first saccade to the top-right, followed by a second saccade to the bottom-left. Right: first saccade to the top-right, followed by a second saccade to the top-left and a third saccade to the bottom-right. Note that the only visual feedback was at the position of the first saccade; additional eye movements were voluntary. Bottom: Directed graph showing transition probability from the first to the second saccade pooled across animals. Left and right nodes correspond to first (task-relevant) and second (voluntary) saccade, respectively. The four exits are arranged from most likely to least likely under an ideal observer model (see Methods) using each animal’s fitted *w_m_*. Arrow width is proportional to transition probability given that a second saccade was made. Results are from manually annotated eye tracking data –– one session per animal (Monkey F: 800 trials; Monkey N: 500 trials). Animals made a second voluntary saccade on more than half the trials (Monkey F: 52.62% of trials; Monkey N: 67.60% of trials). Second saccades were made toward the remaining highest probability exit. Brackets on the right indicate pairwise comparisons between transition probabilities, conditioned on the first saccade (asterisk: statistically significant, two-sided exact binomial test, Bonferroni-corrected, p<0.05).

**Supplementary Figure 2.**
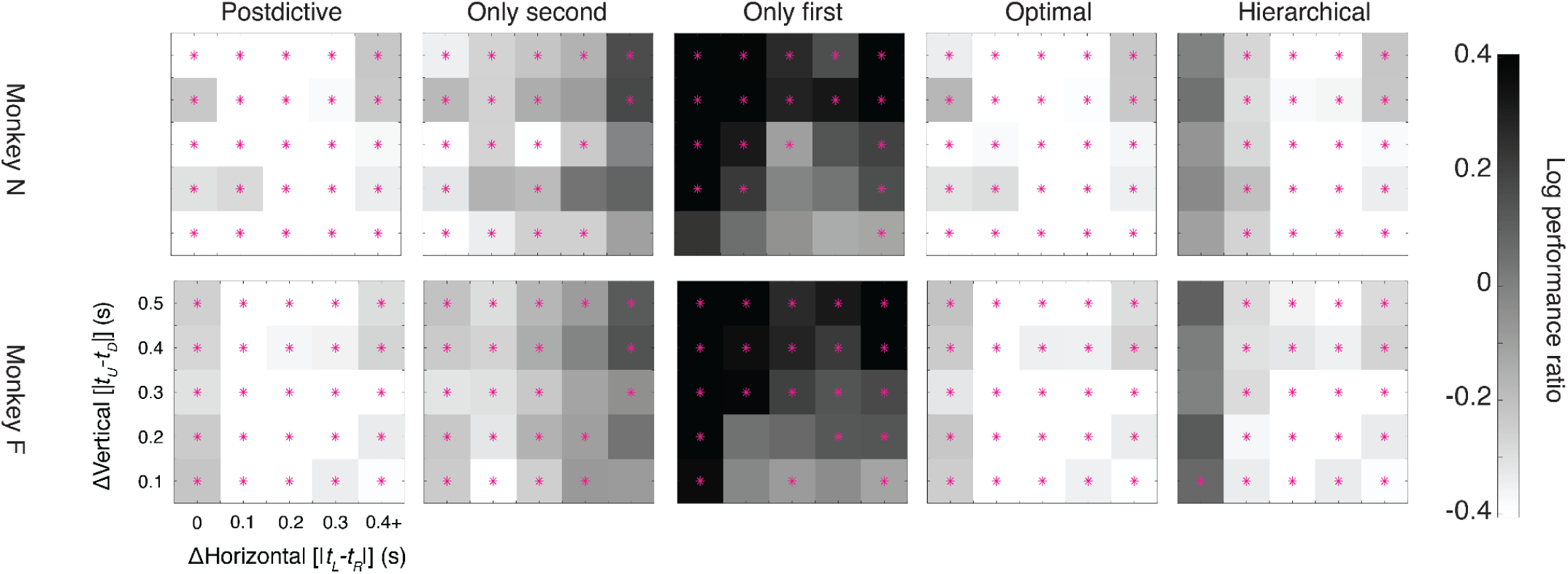
Comparison of monkeys’ choice patterns with several fixed-strategy models. Log performance ratio of animals to fixed strategy models with the same format as Fig. 1j. Asterisks (*) indicates statistical significance of difference between animals and models (two-tailed t-tests, *p* = 0.05).

**Supplementary Figure 3.**
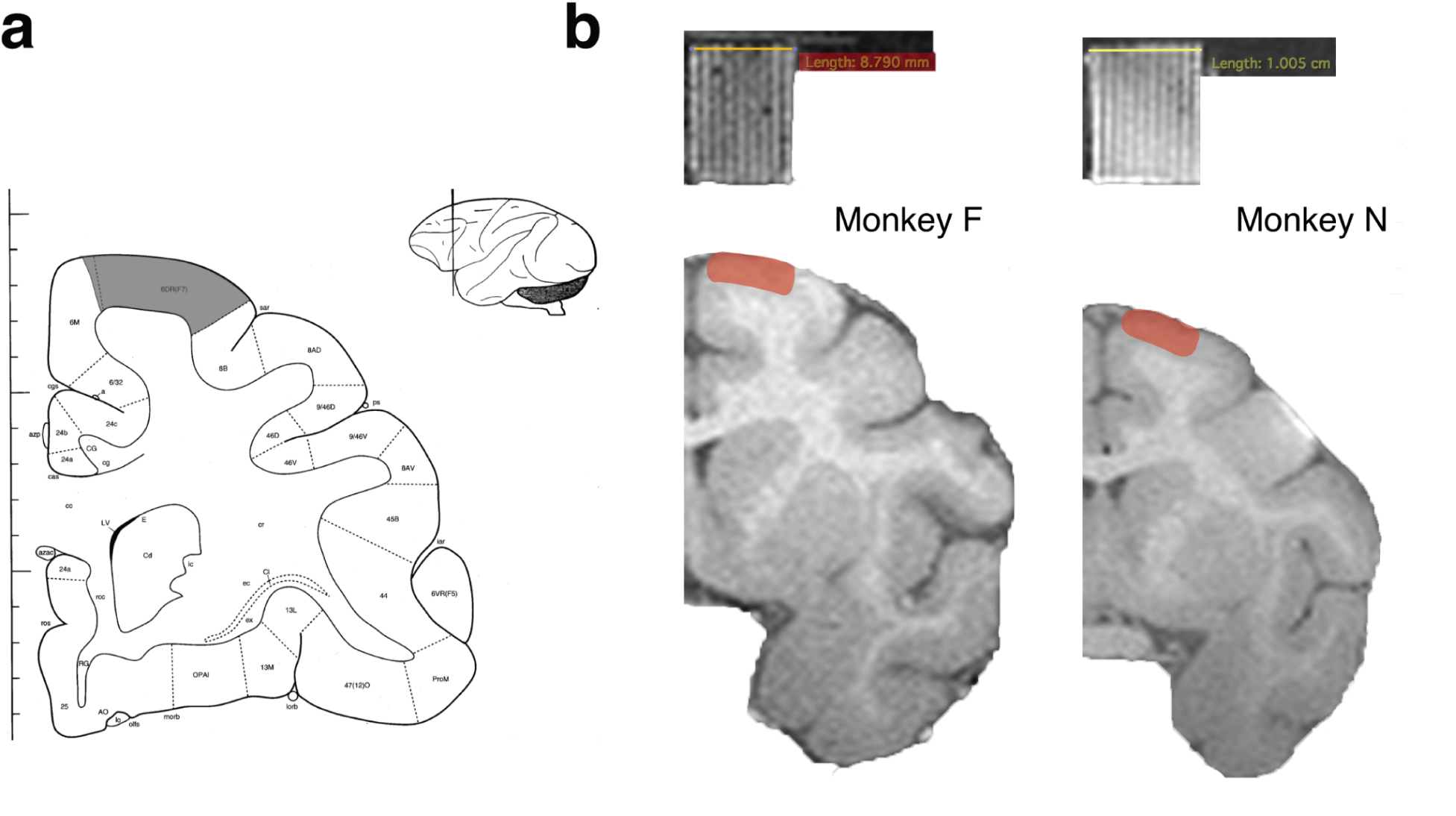
Recording sites. **(a)** Same as Fig. 2a showing schematically the region of interest (gray) over a coronal section. **(b)** Recording sites (red shade) and a recording grid (top grid) shown over a coronal section of the animals’ brain extracted from MRI structural scans. We used the recording grid filled with gadolinium-infused agar gel, which produced bright contrast in MRI, to plan recording trajectories to the targeted brain areas.

**Supplementary Figure 4.**
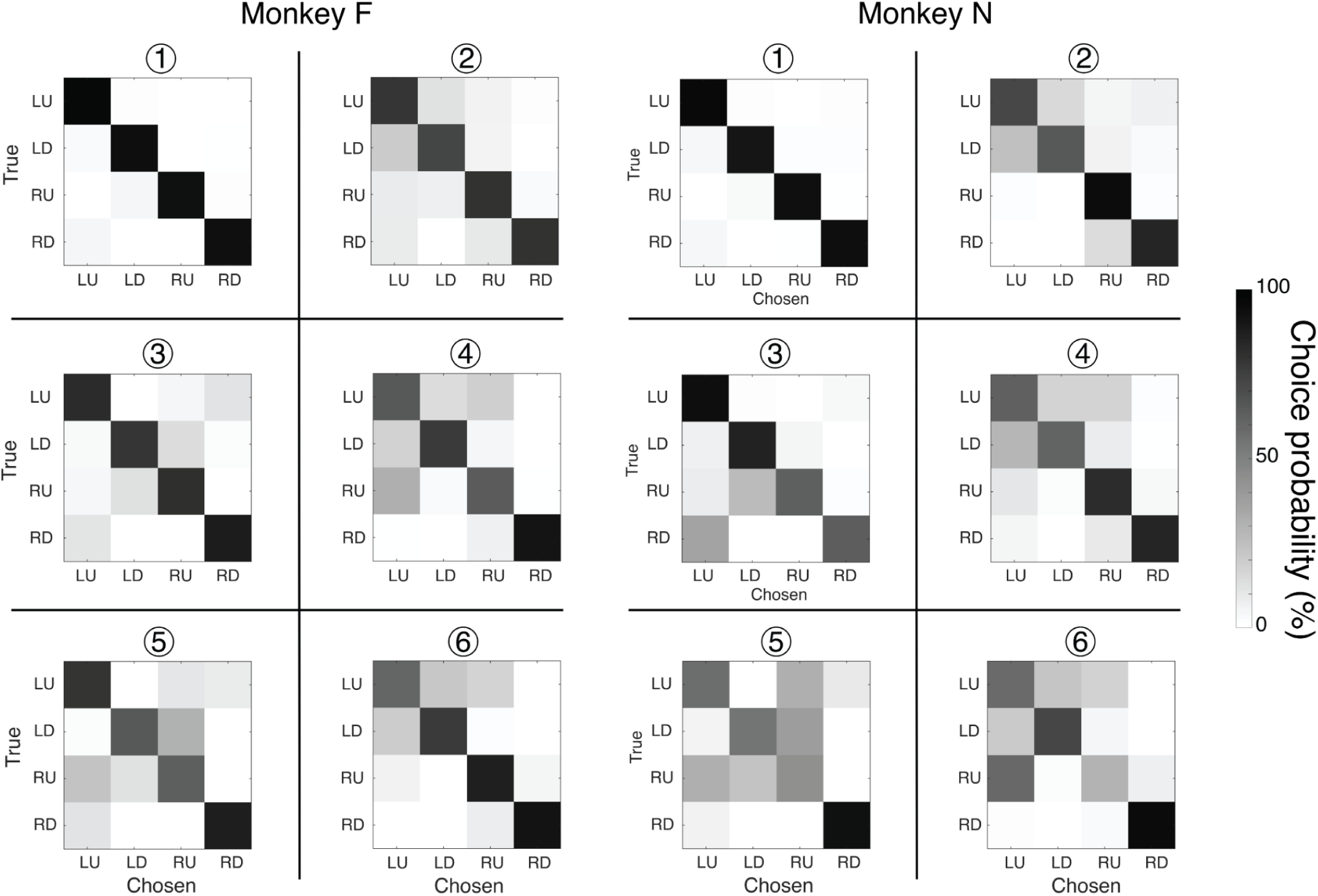
Choice probability as a function of ball exit point for both animals across the six maze geometries used in the physiology experiment. Data were pooled across 27 sessions (14,494 trials) for Monkey F and 34 sessions (22,578 trials) for Monkey N.

**Supplementary Figure 5.**
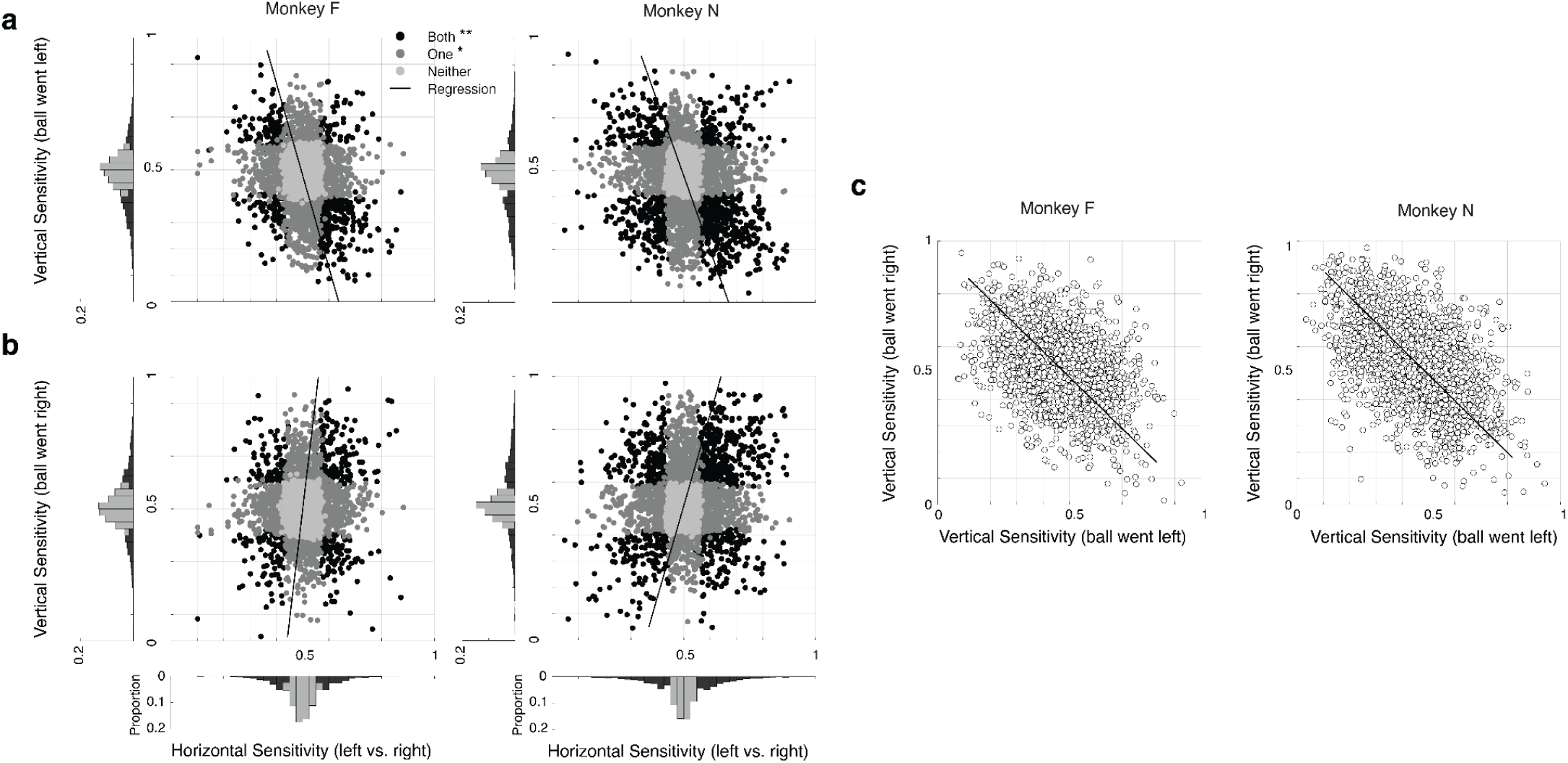
Single neuron sensitivity analysis for the two monkeys. **(a,b)** Scatter plot showing single-neuron sensitivity to flash times associated with the horizontal and vertical arms. Sensitivity was measured using receiver operating characteristic (ROC) using firing rates in 100-ms windows before the relevant flash events (see Methods). The marginals show distribution of sensitivity for horizontal and vertical separately. Results are shown separately for when the ball turned left (a) and right (b). The correlation between horizontal and negative sensitivity was negative when the ball turned left (Monkey F: r=-0.207, p = 1.57e-34; Monkey N: r = r −0.17, p = 3.52e-26), and positive when the ball turned right (Monkey F: r= 0.078, p = 5.093e-06; Monkey N: r = 0.128, p = 2.16e-15). **(c)** Vertical sensitivity compared between trials when the ball moved leftward versus rightward. The two are negatively correlated (Monkey F: r= −0.386; monkey; Monkey N: r=-0.515; both p-values < 1e-80).

**Supplementary Figure 6.**
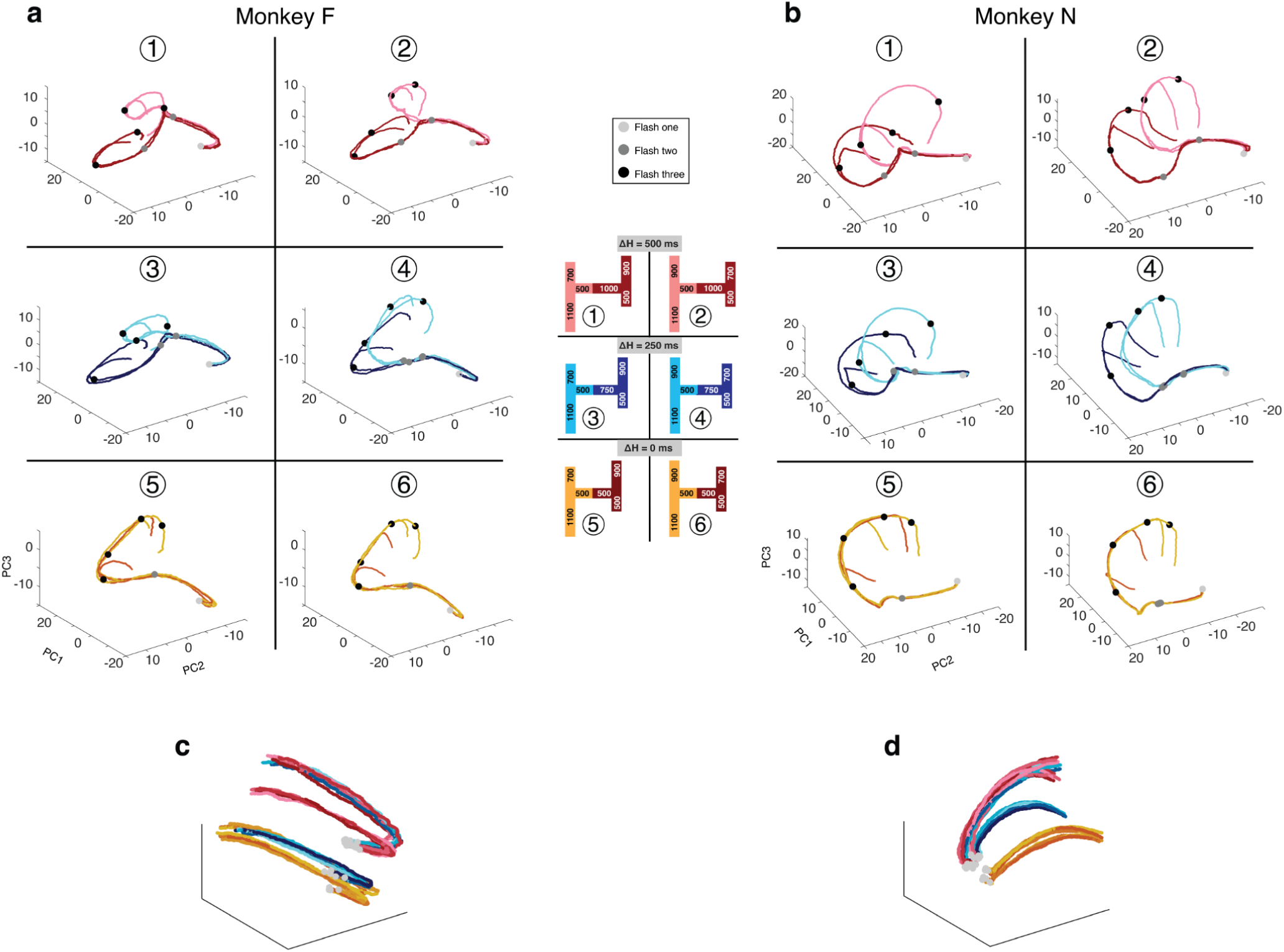
Principal component analysis (PCA) of neural population dynamics. **(a)** Neural trajectories for monkey F from the time of flash 1 to 300 ms after flash 3, shown in the subspace spanned by the top three principal components (PCs), for each of the four possible paths within each maze, using the same color scheme as in the center panel. Neural trajectories from all mazes were projected onto the same PCs and rotated identically for visualization. Both correct and error trials are included. Neural data were pooled across all neurophysiology sessions from monkey F (see Methods). **(b)** Same as in **a**, but for monkey N. **(c)** Same PCA trajectories for monkey F, but with trajectories from all mazes and paths plotted together to highlight clustering of initial conditions and early dynamics from flash 1 to 500 ms after flash 1, before any informative flash cues about ball direction. This interval highlights the two groups of mazes. **(d)** Same as in **c**, but for monkey N.

**Supplementary Figure 7.**
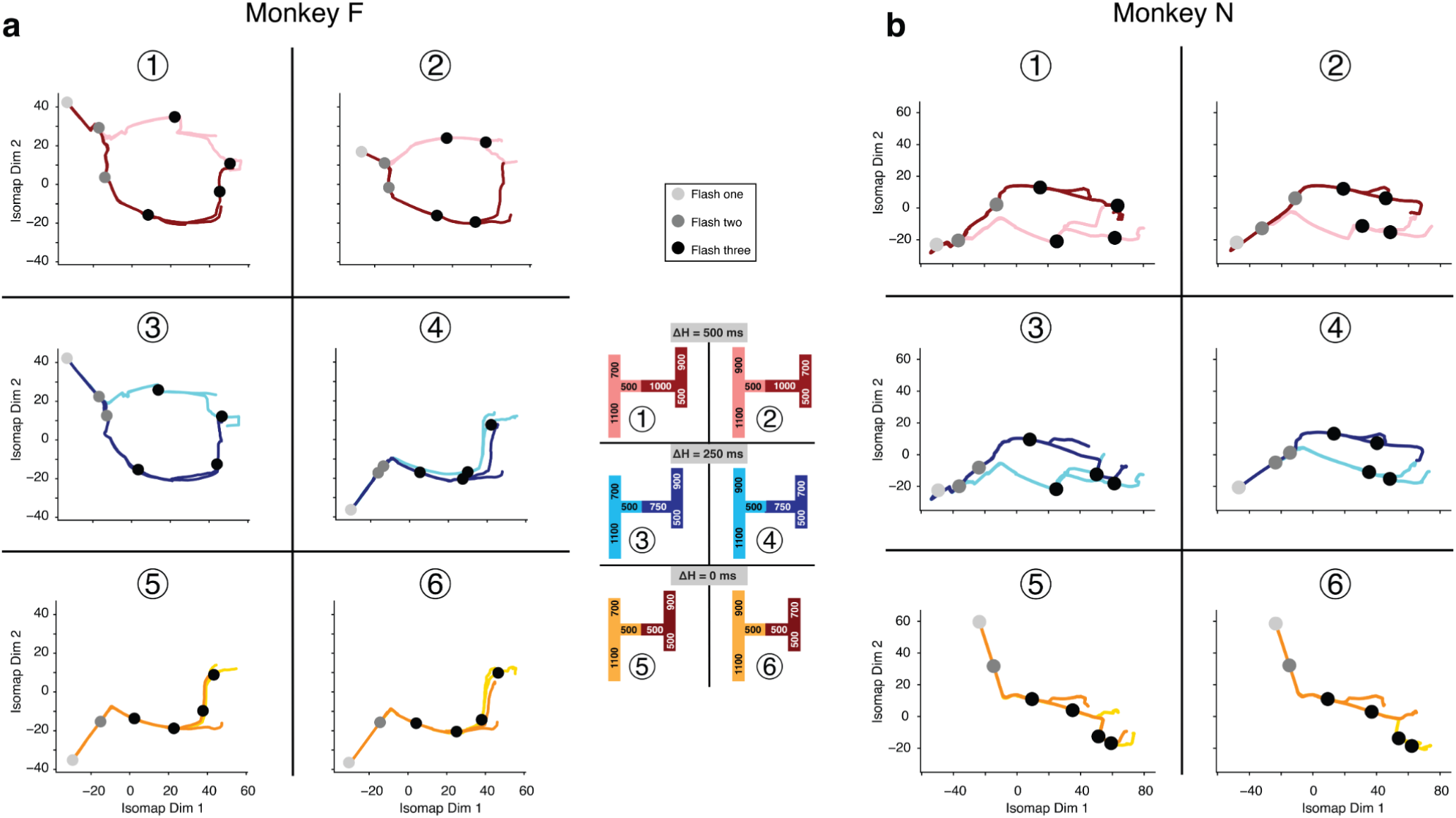
Isomap of neural population dynamics. **(a)** Neural trajectories for monkey F from the time of flash 1 to 300 ms after flash 3, shown in the subspace spanned by the top two Isomap dimensions, for each of the four possible paths within each maze, using the same color scheme as in the center panel. The Isomap embedding was computed on pooled data from all mazes and paths to enable direct comparison across mazes (see methods). Both correct and error trials are included. Neural data were pooled across all neurophysiology sessions from monkey F. **(b)** Same as in **a**, but for monkey N. The center panel is the same as in Supplementary Fig. 6.

**Supplementary Figure 8.**
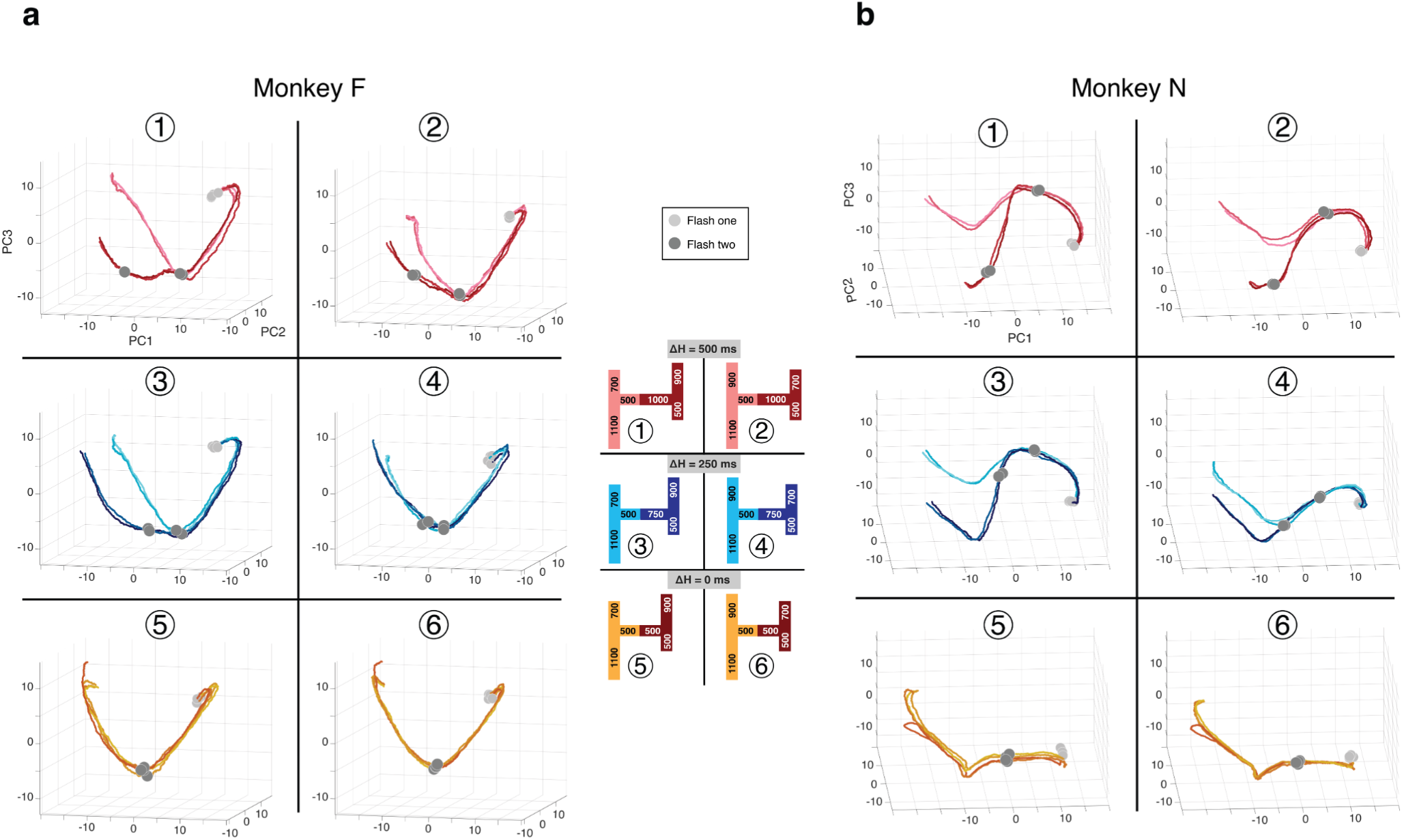
PCA of neural population dynamics conditioned on animal choice. **(a)** Neural trajectories for monkey F from flash 1 to 1,250 ms after flash 1, shown in the subspace spanned by the top three principal components (PCs), for each of the four possible paths within each maze, using the same color scheme as in the center panel. For each path condition, trials were first subdivided according to the animal’s behavioral choice, and neural activity was then averaged separately for left and right choices. The trajectories shown here include only choice-consistent activity, defined as trials in which the animal’s left–right choice matched the true left–right direction of the path. All trajectories from all mazes were projected onto the same PCs and rotated identically for visualization. Neural data were pooled across all neurophysiology sessions from monkey F (see methods). **(b)** Same as in **a**, but for monkey N. The center panel is the same as in Supplementary Fig. 6.

**Supplementary Figure 9.**
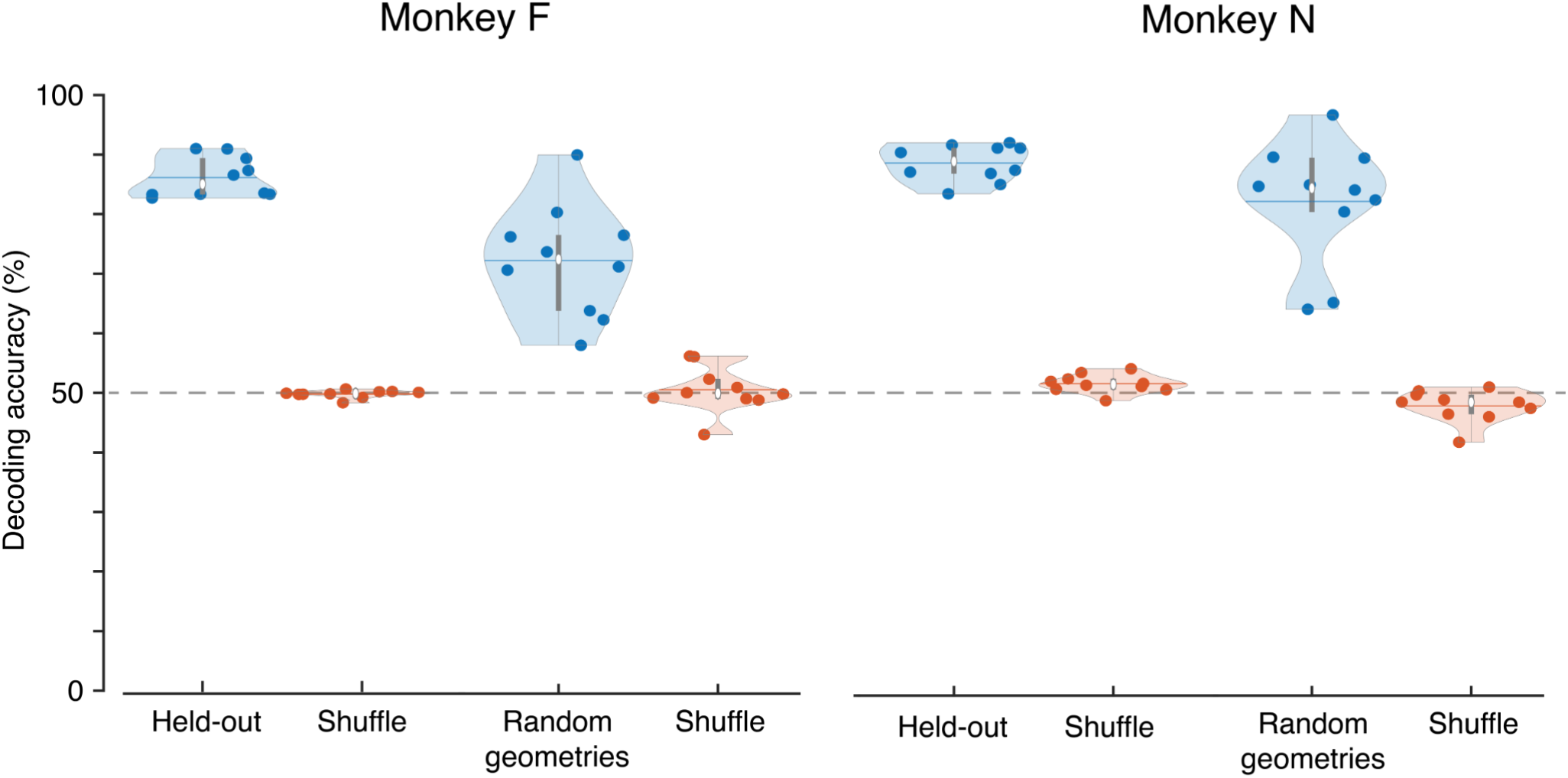
Decoding performance on held-out trials and out-of-distribution random geometries. Decoders were trained separately for each session to predict the animal’s final left–right choice from DMFC population activity, and the y axis shows left–right decoding accuracy. Using repeated 50% holdout splits, class balancing was performed within each training set by subsampling the majority class. “Held-out” denotes performance on held-out test trials, averaged across 200 repeated holdout iterations (see Methods). “Shuffle” denotes the same analysis with randomly permuted training labels to estimate chance performance. “Random geometries” denote performance on the 10% of trials in each session in which mazes with randomly sampled arm lengths were presented; these trials were excluded from both training and held-out test sets and instead used as an out-of-distribution validation set. For this analysis, decoders trained on the standard maze set were evaluated on the random-geometry trials, and performance was averaged across 10 repeated resampling iterations within each session. Each dot represents one session, violins show the distribution across the 10 highest-performing sessions from each animal, white circles indicate the median, horizontal lines indicate the mean, and grey boxes indicate the interquartile range. The dashed horizontal line indicates chance performance.

**Supplementary Figure 10.**
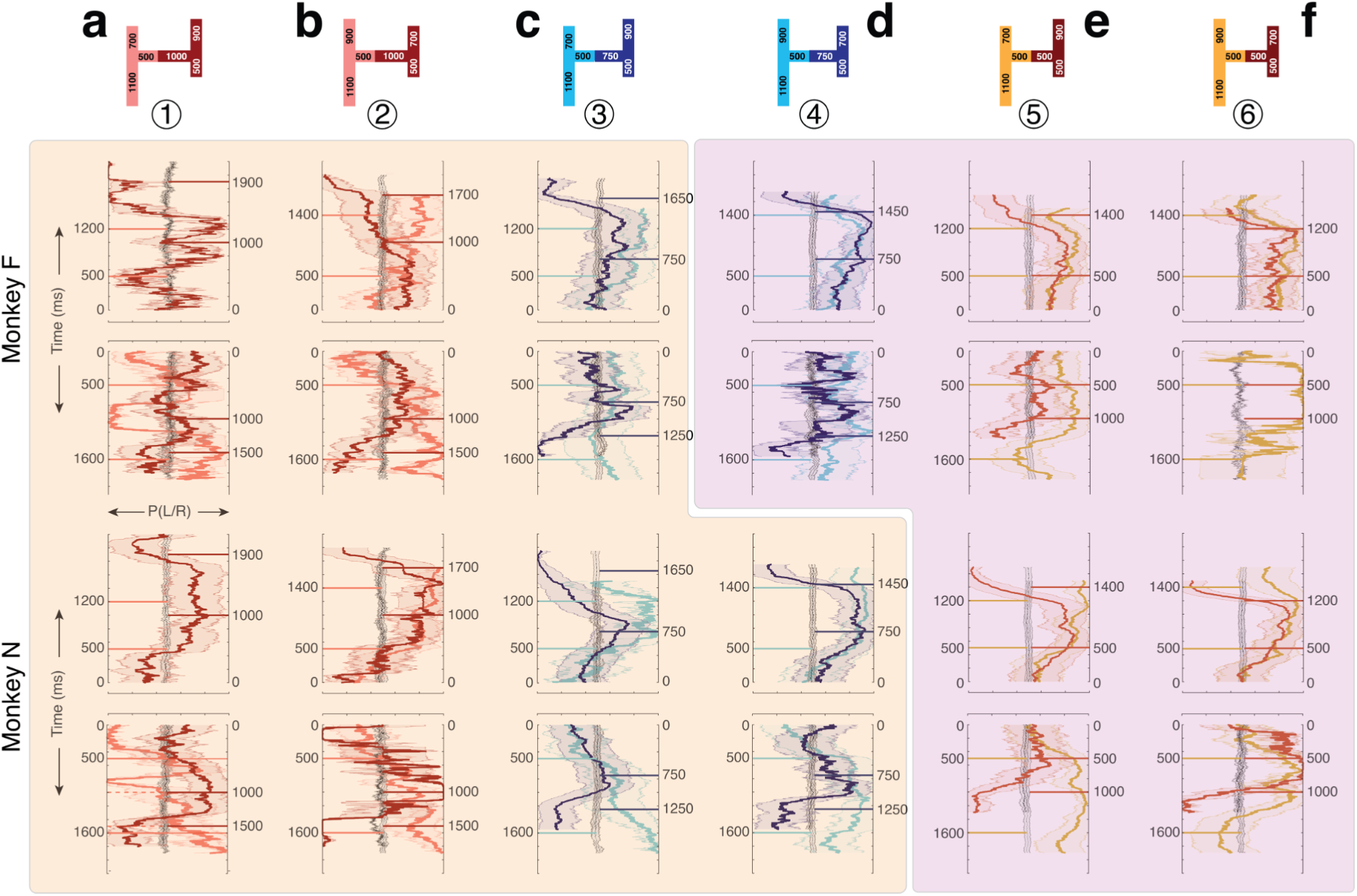
Left–right decision variable (DV) dynamics for error trials. Results are shown separately for the two animals (monkey F: top; monkey N: bottom) across six maze geometries (columns a–f). Results are shown for the 10 highest-performing decoding sessions from each animal (see Methods). For each maze, DV trajectories for the four possible exits are shown in two panels (top panel: left-up and right-up exits, bottom panel: left-down and right-down). The DV (x-axis) is plotted in terms of probability of choosing right (left = 0, right = 1, chance = 0.5) as a function of time (y-axis) from the first flash until 300 ms after the third flash. Time increases upward/downwartd for top/bottom exits. In each panel, mean DV (solid lines) ± 0.5 standard deviations across trials (shaded areas) are color-matched to maze paths (top). Each panel also shows trial-shuffled mean DV (black) ± 0.5 standard deviations across trials (gray). Horizontal solid lines mark the timing of the second and third flashes for each exit. Background shadings separate mazes based on strategy inferred based on decoding results from all trials, same as in figure 3 (yellow: hierarchical, magenta: sequential).

**Supplementary Figure 11.**
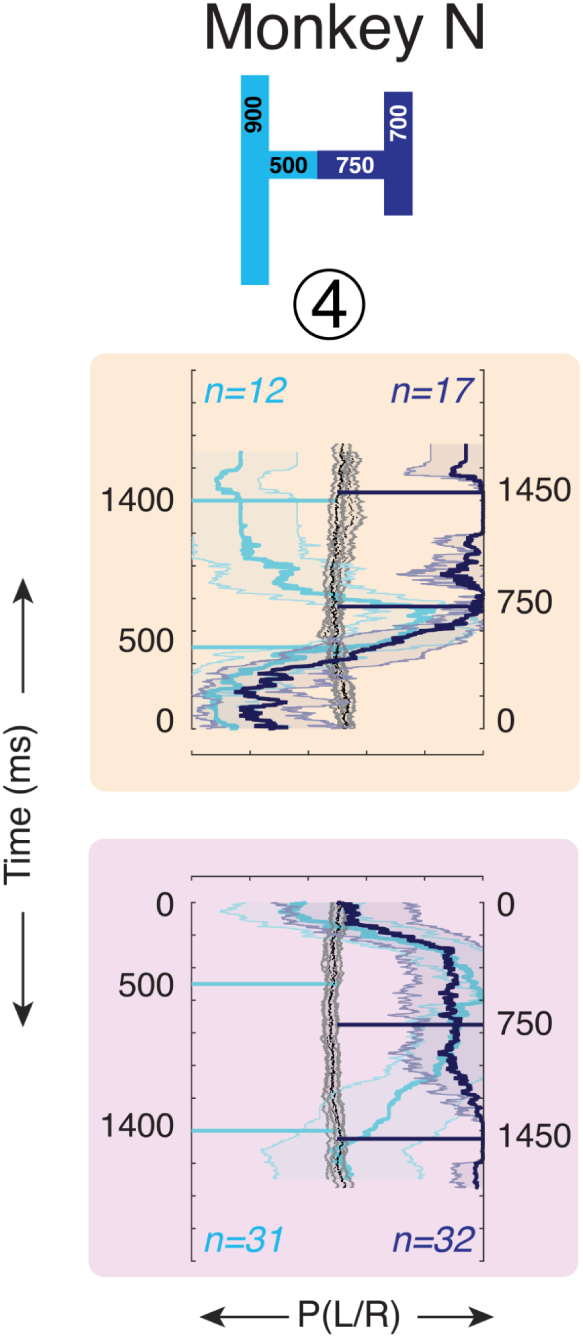
Left–right decision variable (DV) for the upper exits in maze 4 for monkey N. Trials are grouped by neurally inferred initial-condition state (top, hierarchical; bottom, sequential), defined from single-trial population activity measured from flash 1 to 500 ms after flash 1 using hierarchical clustering, and plotted in the same format as Fig. 3. In each panel, mean DV (solid lines) ± 0.5 standard deviations across trials (shaded areas) are color-matched to maze paths (top). Each panel also shows trial-shuffled mean DV (black) ± 0.5 standard deviations across trials (gray). Horizontal solid lines mark the timing of the second and third flashes for each exit. Background shadings separate mazes based on strategy inferred based on decoding results from all trials, same as in figure 3 (yellow: hierarchical, magenta: sequential).The numbers at each corner indicate the number of trials associated with each exit. The decoder was trained to predict the monkey’s realized final left-versus-right choice from endpoint population activity measured from flash 3 to 300 ms after flash 3 (see methods).

---

**Table S1.**
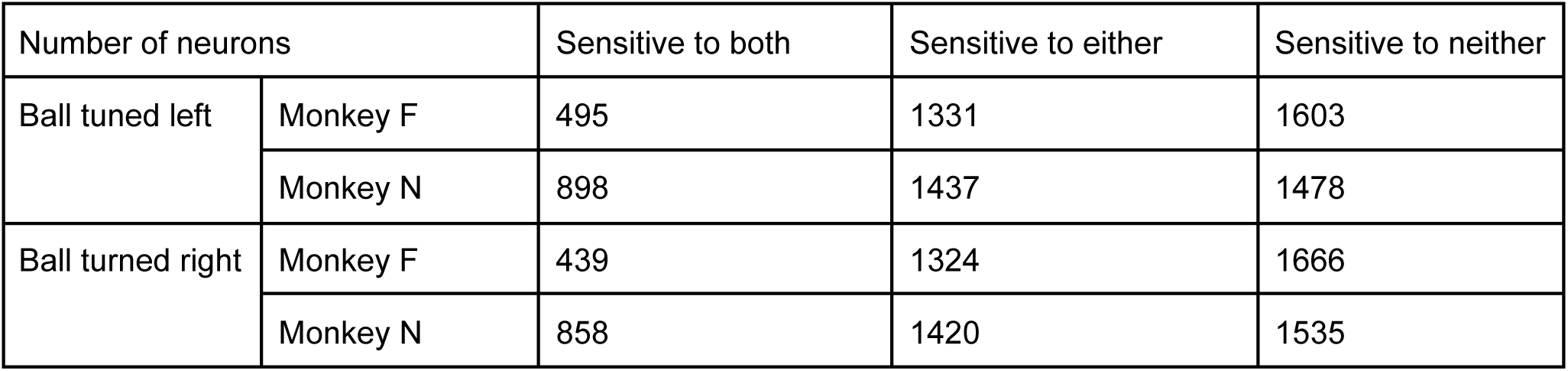
Number of neurons with significant sensitivity to the flash times associated with discriminating both horizontal and vertical arms (3rd column), either horizontal or vertical arms but not both (4th column), and neither (5th column).

## Notes

### Competing Interest Statement

The authors have declared no competing interest.

## References

1. Newell, A. & Simon, H. A. Human Problem Solving. (Echo Point Books & Media, 2023).

2. Tversky, A. & Kahneman, D. Judgment under uncertainty: Heuristics and biases. in Handbook of the Fundamentals of Financial Decision Making 261–268 (WORLD SCIENTIFIC, 2013).

3. Johnson-Laird, P. N. Kahneman, Tversky and Kahneman-Tversky: three ways of thinking. Think. Reason. 30, 531–547 (2024).

4. Gigerenzer, G., Hertwig, R. & Pachur, T. Heuristics. (Oxford University Press, New York, NY, 2016).

5. Gentner, D. & Smith, L. A. Analogical Learning and Reasoning. (Oxford University Press, 2013).

6. Morrison, R. G. & Holyoak, K. J. Problem Solving. in Encyclopedia of the Neurological Sciences 60–63 (Elsevier, 2003).

7. Miller, E. K. & Cohen, J. D. An integrative theory of prefrontal cortex function. Annu. Rev. Neurosci. 24, 167–202 (2001).

8. Ho, M. K. et al. People construct simplified mental representations to plan. Nature 606, 129–136 (2022).

9. Dayan, P. & Daw, N. D. Decision theory, reinforcement learning, and the brain. Cogn. Affect. Behav. Neurosci. 8, 429–453 (2008).

10. Huys, Q. J. M. et al. Interplay of approximate planning strategies. Proc. Natl. Acad. Sci. U. S. A. 112, 3098–3103 (2015).

11. Lake, B. M., Ullman, T. D., Tenenbaum, J. B. & Gershman, S. J. Building machines that learn and think like people. Behav. Brain Sci. 40, e253 (2017).

12. Griffiths, T. L., Kemp, C. & Tenenbaum, J. B. Bayesian models of cognition. in The Cambridge Handbook of Computational Cognitive Sciences 80–138 (Cambridge University Press, 2023).

13. Eckstein, M. K. & Collins, A. G. E. How the mind creates structure: Hierarchical learning of action sequences. CogSci 43, 618–624 (2021).

14. Heald, J. B., Lengyel, M. & Wolpert, D. M. Contextual inference underlies the learning of sensorimotor repertoires. Nature 600, 489–493 (2021).

15. Dehaene, S., Kerszberg, M. & Changeux, J.-P. A neuronal model of a global workspace in effortful cognitive tasks. Ann. N. Y. Acad. Sci. 929, 152–165 (2006).

16. Dehaene, S. & Changeux, J. P. A hierarchical neuronal network for planning behavior. Proc. Natl. Acad. Sci. U. S. A. 94, 13293–13298 (1997).

17. Rougier, N. P., Noelle, D. C., Braver, T. S., Cohen, J. D. & O’Reilly, R. C. Prefrontal cortex and flexible cognitive control: rules without symbols. Proc. Natl. Acad. Sci. U. S. A. 102, 7338–7343 (2005).

18. Shenhav, A., Botvinick, M. M. & Cohen, J. D. The expected value of control: an integrative theory of anterior cingulate cortex function. Neuron 79, 217–240 (2013).

19. Frömer, R. & Shenhav, A. Filling the gaps: Cognitive control as a critical lens for understanding mechanisms of value-based decision-making. Neurosci. Biobehav. Rev. 134, 104483 (2022).

20. Fischer, J., Mikhael, J. G., Tenenbaum, J. B. & Kanwisher, N. Functional neuroanatomy of intuitive physical inference. Proc. Natl. Acad. Sci. U. S. A. 113, E5072–81 (2016).

21. Kroger, J. K. et al. Recruitment of anterior dorsolateral prefrontal cortex in human reasoning: a parametric study of relational complexity. Cereb. Cortex 12, 477–485 (2002).

22. Koechlin, E. & Summerfield, C. An information theoretical approach to prefrontal executive function. Trends Cogn. Sci. 11, 229–235 (2007).

23. Koechlin, E., Basso, G., Pietrini, P., Panzer, S. & Grafman, J. The role of the anterior prefrontal cortex in human cognition. Nature 399, 148–151 (1999).

24. Dehaene, S., Piazza, M., Pinel, P. & Cohen, L. Three parietal circuits for number processing. Cogn. Neuropsychol. 20, 487–506 (2003).

25. Mnih, V. et al. Human-level control through deep reinforcement learning. Nature 518, 529–533 (2015).

26. Silver, D., et al. Mastering chess and shogi by self-play with a general reinforcement learning algorithm. arXiv [cs.AI] (2017).

27. Silver, D. et al. Mastering the game of Go without human knowledge. Nature 550, 354–359 (2017).

28. Brown, T. B. et al. Language Models are Few-Shot Learners. arXiv [cs.CL] (2020).

29. Baker, B., et al. Emergent Tool Use From Multi-Agent Autocurricula. arXiv [cs.LG] (2019).

30. Schulman, J., Wolski, F., Dhariwal, P., Radford, A. & Klimov, O. Proximal Policy Optimization Algorithms. arXiv [cs.LG] (2017).

31. Pearl, J. Heuristics. (Addison Wesley Longman Publishing, New York, NY, 1984).

32. Shadlen, M. N. & Newsome, W. T. Motion perception: seeing and deciding. Proc. Natl. Acad. Sci. U. S. A. 93, 628–633 (1996).

33. de Lafuente, V. & Romo, R. Neuronal correlates of subjective sensory experience. Nat. Neurosci. 8, 1698–1703 (2005).

34. Luna, R., Hernández, A., Brody, C. D. & Romo, R. Neural codes for perceptual discrimination in primary somatosensory cortex. Nat. Neurosci. 8, 1210–1219 (2005).

35. Brunton, B. W., Botvinick, M. M. & Brody, C. D. Rats and Humans Can Optimally Accumulate Evidence for Decision-Making. Science 340, 95–98 (2013).

36. Beck, J. M. et al. Probabilistic population codes for Bayesian decision making. Neuron 60, 1142–1152 (2008).

37. Kiani, R. & Shadlen, M. N. Representation of confidence associated with a decision by neurons in the parietal cortex. Science 324, 759–764 (2009).

38. Nieh, E. H. et al. Geometry of abstract learned knowledge in the hippocampus. Nature 595, 80–84 (2021).

39. Harvey, C. D., Coen, P. & Tank, D. W. Choice-specific sequences in parietal cortex during a virtual-navigation decision task. Nature 484, 62–68 (2012).

40. Drugowitsch, J., DeAngelis, G. C., Klier, E. M., Angelaki, D. E. & Pouget, A. Optimal multisensory decision-making in a reaction-time task. Elife 3, (2014).

41. Churchland, A. K., Kiani, R. & Shadlen, M. N. Decision-making with multiple alternatives. Nat. Neurosci. 11, 693–702 (2008).

42. Licata, A. M. et al. Posterior parietal cortex guides visual decisions in rats. J. Neurosci. 37, 4954–4966 (2017).

43. Rossi-Pool, R. et al. Decoding a Decision Process in the Neuronal Population of Dorsal Premotor Cortex. Neuron 96, 1432–1446.e7 (2017).

44. Platt, M. L. & Glimcher, P. W. Neural correlates of decision variables in parietal cortex. Nature 400, 233–238 (1999).

45. Peixoto, D. et al. Decoding and perturbing decision states in real time. Nature 591, 604–609 (2021).

46. Vivar-Lazo, M. & Fetsch, C. R. Neural basis of concurrent deliberation toward a choice and degree of confidence. bioRxiv (2024) doi:10.1101/2024.08.06.606833.

47. Boundy-Singer, Z. M., Ziemba, C. M. & Goris, R. L. T. Sensory population activity reveals confidence computations in the primate visual system. bioRxiv (2024) doi:10.1101/2024.08.01.606172.

48. Ghose, G. M. & Maunsell, J. H. R. Attentional modulation in visual cortex depends on task timing. Nature 419, 616–620 (2002).

49. Cohen, M. R. & Maunsell, J. H. R. Attention improves performance primarily by reducing interneuronal correlations. Nat. Neurosci. 12, 1594–1600 (2009).

50. Seidemann, E., Zohary, E. & Newsome, W. T. Temporal gating of neural signals during performance of a visual discrimination task. Nature 394, 72–75 (1998).

51. Zylberberg, A., Fetsch, C. R. & Shadlen, M. N. The influence of evidence volatility on choice, reaction time and confidence in a perceptual decision. Elife 5, (2016).

52. Cisek, P. Making decisions through a distributed consensus. Curr. Opin. Neurobiol. 22, 927–936 (2012).

53. Glaze, C. M., Kable, J. W. & Gold, J. I. Normative evidence accumulation in unpredictable environments. Elife 4, (2015).

54. Castiñeiras, J. R. & Renart, A. Control Limited Perceptual Decision Making. bioRxiv 2022.06.24.497481 (2022) doi:10.1101/2022.06.24.497481.

55. Okazawa, G., Hatch, C. E., Mancoo, A., Machens, C. K. & Kiani, R. Representational geometry of perceptual decisions in the monkey parietal cortex. Cell 184, 3748–3761.e18 (2021).

56. Lee, D., McGreevy, B. P. & Barraclough, D. J. Learning and decision making in monkeys during a rock–paper–scissors game. Cognitive Brain Research 25, 416–430 (2005).

57. Angelaki, D. E., Gu, Y. & Deangelis, G. C. Visual and vestibular cue integration for heading perception in extrastriate visual cortex. J. Physiol. 589, 825–833 (2011).

58. Graf, A. B. A., Kohn, A., Jazayeri, M. & Movshon, J. A. Decoding the activity of neuronal populations in macaque primary visual cortex. Nat. Neurosci. 14, 239–245 (2011).

59. Stine, G. M., Trautmann, E. M., Jeurissen, D. & Shadlen, M. N. A neural mechanism for terminating decisions. Neuron 111, 2601–2613.e5 (2023).

60. International Brain Laboratory et al. Correction: Standardized and reproducible measurement of decision-making in mice. Elife 11, (2022).

61. Mante, V., Sussillo, D., Shenoy, K. V. & Newsome, W. T. Context-dependent computation by recurrent dynamics in prefrontal cortex. Nature 503, 78–84 (2013).

62. Sohn, H., Narain, D., Meirhaeghe, N. & Jazayeri, M. Bayesian computation through cortical latent dynamics. Neuron 103, 934–947.e5 (2019).

63. Egger, S. W., Remington, E. D., Chang, C.-J. & Jazayeri, M. Internal models of sensorimotor integration regulate cortical dynamics. Nat. Neurosci. 22, 1871–1882 (2019).

64. Wang, J., Narain, D., Hosseini, E. A. & Jazayeri, M. Flexible timing by temporal scaling of cortical responses. Nat. Neurosci. 21, 102–110 (2018).

65. Remington, E. D., Narain, D., Hosseini, E. A. & Jazayeri, M. Flexible Sensorimotor Computations through Rapid Reconfiguration of Cortical Dynamics. Neuron 98, 1005–1019.e5 (2018).

66. Driscoll, L. N., Shenoy, K. & Sussillo, D. Flexible multitask computation in recurrent networks utilizes shared dynamical motifs. Nat. Neurosci. 27, 1349–1363 (2024).

67. Sarafyazd, M. & Jazayeri, M. Hierarchical reasoning by neural circuits in the frontal cortex. Science 364, (2019).

68. Xue, C., Kramer, L. E. & Cohen, M. R. Dynamic task-belief is an integral part of decision-making. Neuron 110, 2503–2511.e3 (2022).

69. Xue, C., Markman, S. K., Chen, R., Kramer, L. E. & Cohen, M. R. Task interference as a neuronal basis for the cost of cognitive flexibility. bioRxiv (2024) doi:10.1101/2024.03.04.583375.

70. Nassar, M. R., Wilson, R. C., Heasly, B. & Gold, J. I. An approximately Bayesian delta-rule model explains the dynamics of belief updating in a changing environment. J. Neurosci. 30, 12366–12378 (2010).

71. Li, Y. S., Nassar, M. R., Kable, J. W. & Gold, J. I. Individual neurons in the cingulate cortex encode action monitoring, not selection, during adaptive decision-making. J. Neurosci. 39, 6668–6683 (2019).

72. Wang, J., Hosseini, E., Meirhaeghe, N., Akkad, A. & Jazayeri, M. Reinforcement regulates timing variability in thalamus. Elife 9, (2020).

73. Beiran, M., Meirhaeghe, N., Sohn, H., Jazayeri, M. & Ostojic, S. Parametric control of flexible timing through low-dimensional neural manifolds. Neuron (2023) doi:10.1016/j.neuron.2022.12.016.

74. Meirhaeghe, N., Sohn, H. & Jazayeri, M. A precise and adaptive neural mechanism for predictive temporal processing in the frontal cortex. Neuron 109, 2995–3011.e5 (2021).

75. Cadena-Valencia, J., García-Garibay, O., Merchant, H., Jazayeri, M. & de Lafuente, V. Entrainment and maintenance of an internal metronome in supplementary motor area. Elife 7, (2018).

76. Merchant, H., Harrington, D. L. & Meck, W. H. Neural basis of the perception and estimation of time. Annu. Rev. Neurosci. 36, 313–336 (2013).

77. Merchant, H., Perez, O., Zarco, W. & Gamez, J. Interval Tuning in the Primate Medial Premotor Cortex as a General Timing Mechanism. J. Neurosci. 33, 9082–9096 (2013).

78. Emmons, E. B. et al. Rodent Medial Frontal Control of Temporal Processing in the Dorsomedial Striatum. J. Neurosci. 37, 8718–8733 (2017).

79. Murakami, M., Shteingart, H., Loewenstein, Y. & Mainen, Z. F. Distinct Sources of Deterministic and Stochastic Components of Action Timing Decisions in Rodent Frontal Cortex. Neuron 94, 908–919.e7 (2017).

80. Mita, A., Mushiake, H., Shima, K., Matsuzaka, Y. & Tanji, J. Interval time coding by neurons in the presupplementary and supplementary motor areas. Nat. Neurosci. 12, 502–507 (2009).

81. Gibbon, J. & Church, R. M. Representation of time. Cognition 37, 23–54 (1990).

82. Ramadan, M., Tang, C., Watters, N. & Jazayeri, M. Computational basis of hierarchical and counterfactual information processing. Nature Human Behaviour 1–15 (2025).

83. Rajalingham, R., Sohn, H. & Jazayeri, M. Dynamic tracking of objects in the macaque dorsomedial frontal cortex. Nat. Commun. 16, 346 (2025).

84. Jun, J. J. et al. Fully integrated silicon probes for high-density recording of neural activity. Nature 551, 232–236 (2017).

85. Trautmann, E. M. et al. Large-scale high-density brain-wide neural recording in nonhuman primates. Nat. Neurosci. 28, 1562–1575 (2025).

86. Hanks, T. D. et al. Distinct relationships of parietal and prefrontal cortices to evidence accumulation. Nature 520, 220–223 (2015).

87. Gosztolai, A., Peach, R. L., Arnaudon, A., Barahona, M. & Vandergheynst, P. MARBLE: interpretable representations of neural population dynamics using geometric deep learning. Nat. Methods 1–9 (2025).

88. Padoa-Schioppa, C. & Assad, J. A. Neurons in the orbitofrontal cortex encode economic value. Nature 441, 223–226 (2006).

89. Kepecs, A., Uchida, N., Zariwala, H. A. & Mainen, Z. F. Neural correlates, computation and behavioural impact of decision confidence. Nature 455, 227–231 (2008).

90. Li, N., Chen, T.-W., Guo, Z. V., Gerfen, C. R. & Svoboda, K. A motor cortex circuit for motor planning and movement. Nature vol. 519 51–56 Preprint at 10.1038/nature14178 (2015).

91. Nogueira, R., Esteki, S., Fusi, S. & Kiani, R. The geometry of context-dependent biased decisions during learning. bioRxivorg (2026) doi:10.64898/2026.01.14.699497.

92. Chen, R., Radkani, S., Valluru, N., Yoo, S. B. M. & Jazayeri, M. Evidence accumulation from experience and observation in the cingulate cortex. Nature 1–9 (2026).

93. Wallis, J. D., Anderson, K. C. & Miller, E. K. Single neurons in prefrontal cortex encode abstract rules. Nature 411, 953–956 (2001).

94. Pagan, M. et al. Individual variability of neural computations underlying flexible decisions. Nature 639, 421–429 (2025).

95. Wimmer, R. D. et al. Thalamic control of sensory selection in divided attention. Nature 526, 705–709 (2015).

96. International Brain Laboratory et al. A brain-wide map of neural activity during complex behaviour. Nature 645, 177–191 (2025).

97. Barraclough, D. J., Conroy, M. L. & Lee, D. Prefrontal cortex and decision making in a mixed-strategy game. Nat. Neurosci. 7, 404–410 (2004).

98. Yang, Q. et al. Monkey plays Pac-Man with compositional strategies and hierarchical decision-making. Elife 11, (2022).

99. Callaway, F. et al. Rational use of cognitive resources in human planning. *Nat*. Hum. Behav. 6, 1112–1125 (2022).

100. Zylberberg, A. Decision prioritization and causal reasoning in decision hierarchies. PLoS Comput. Biol. 17, e1009688 (2021).

101. Boureau, Y.-L., Sokol-Hessner, P. & Daw, N. D. Deciding how to decide: Self-control and meta-decision making. Trends Cogn. Sci. 19, 700–710 (2015).

102. Krakauer, J. W., Ghazanfar, A. A., Gomez-Marin, A., MacIver, M. A. & Poeppel, D. Neuroscience Needs Behavior: Correcting a Reductionist Bias. Neuron 93, 480–490 (2017).

103. Shenoy, K. V., Sahani, M. & Churchland, M. M. Cortical control of arm movements: a dynamical systems perspective. Annu. Rev. Neurosci. 36, 337–359 (2013).

104. Sohn, H., Meirhaeghe, N., Rajalingham, R. & Jazayeri, M. A Network Perspective on Sensorimotor Learning. Trends Neurosci. 44, 170–181 (2021).

105. Remington, E. D., Egger, S. W., Narain, D., Wang, J. & Jazayeri, M. A Dynamical Systems Perspective on Flexible Motor Timing. Trends Cogn. Sci. 22, 938–952 (2018).

106. Vyas, S., Golub, M. D., Sussillo, D. & Shenoy, K. V. Computation Through Neural Population Dynamics. Annu. Rev. Neurosci. 43, 249–275 (2020).

107. Ashwood, Z. C. et al. Mice alternate between discrete strategies during perceptual decision-making. Nat. Neurosci. 25, 201–212 (2022).

108. Le, N. M. et al. Mixtures of strategies underlie rodent behavior during reversal learning. PLoS Comput. Biol. 19, e1011430 (2023).

109. Mohammadi, Z., Ashwood, Z. C., International Brain Laboratory & Pillow, J. W. Identifying the factors governing internal state switches during nonstationary sensory decision-making. Nat. Commun. 16, 11684 (2025).

110. Botvinick, M. M., Braver, T. S., Barch, D. M., Carter, C. S. & Cohen, J. D. Conflict monitoring and cognitive control. Psychol. Rev. 108, 624–652 (2001).

111. Collins, A. G. E. & Frank, M. J. Cognitive control over learning: creating, clustering, and generalizing task-set structure. Psychol. Rev. 120, 190–229 (2013).

112. Lieder, F. & Griffiths, T. L. Resource-rational analysis: Understanding human cognition as the optimal use of limited computational resources. Behav. Brain Sci. 43, e1 (2019).

113. Gibson, J. J. The Ecological Approach to Visual Perception: Classic Edition. (Psychology Press, 2014).

114. Malapani, C. & Fairhurst, S. Scalar Timing in Animals and Humans. Learn. Motiv. 33, 156–176 (2002).

115. Green, D. M. & Swets, J. A. Signal detection theory and psychophysics. 455, (1966).

116. Sohn, H., Narain, D., Meirhaeghe, N. & Jazayeri, M. Bayesian Computation through Cortical Latent Dynamics. Neuron 103, 934–947.e5 (2019).

117. Fujii, N., Mushiake, H. & Tanji, J. Distribution of eye- and arm-movement-related neuronal activity in the SEF and in the SMA and Pre-SMA of monkeys. J. Neurophysiol. 87, 2158–2166 (2002).

118. Huerta, M. F. & Kaas, J. H. Supplementary eye field as defined by intracortical microstimulation: connections in macaques. J. Comp. Neurol. 293, 299–330 (1990).

119. Matsuzaka, Y., Aizawa, H. & Tanji, J. A motor area rostral to the supplementary motor area (presupplementary motor area) in the monkey: neuronal activity during a learned motor task. J. Neurophysiol. 68, 653–662 (1992).

120. Ito, S., Stuphorn, V., Brown, J. W. & Schall, J. D. Performance monitoring by the anterior cingulate cortex during saccade countermanding. Science 302, 120–122 (2003).

121. Arya, S., Mount, D. M., Netanyahu, N. S., Silverman, R. & Wu, A. Y. An optimal algorithm for approximate nearest neighbor searching fixed dimensions. J. ACM 45, 891–923 (1998).

122. Berry, T. & Sauer, T. Consistent manifold representation for topological data analysis. Found. Data Sci. 1, 1–38 (2019).

